# Curcumin alleviates Arsenic trioxide-induced behavioural impairment, oxidative damage & morphological alterations in striatal region of mice brain

**DOI:** 10.1101/2023.12.14.571716

**Authors:** Kamlesh Kumar Pandey, Kamakshi Mehta, Balpreet Kaur, Pushpa Dhar, Saroj Kaler

**Affiliations:** Department of Anatomy, All India Institute of Medical Sciences, New Delhi-110029

## Abstract

Arsenic-induced neurotoxicity is well-documented in literature and reported to have dose-dependent damaging effects in the mice brain. Curcumin, a cost-effective plant polyphenol, safely demonstrates protective effects against arsenic-induced neurotoxicity by modifying oxidative stress, apoptosis, and neurochemistry in rodents’ brain.

The present study determined the neuroprotective potential of curcumin (CUR) on adverse effects induced by arsenic trioxide (As_2_O_3_) in mice striatal region. Healthy adult male mice were chronically administered with varying concentrations of As_2_O_3_ (2, 4 & 8 mg/kg bw) alone and along with CUR (100 mg/kg bw) via oral route for 45 days. Towards the end of experimental period, the animals were subjected to behavioural paradigm including open field task, novel object recognition, rota-rod and morris water maze. Fresh striatal tissues were collected from the animals on day 46 for biochemical analysis such as MDA, GPx and GSH. While perfusion fixed brains were processed for morphological observations.

Behavioural study showed an apparent decrease in certain cognitive functions (learning and memory) and locomotor activity in mice exposed to As_2_O_3_ compared to controls. Simultaneous treatment of As_2_O_3_ (2, 4 & 8 mg/kg bw) and curcumin (100 mg/kg bw) alleviated the *As*-induced locomotor and cognitive deficits. As_2_O_3_ alone exposure also exhibited a significant increase in oxidative stress marker (MDA) and decrease in antioxidant enzyme levels (GPx, GSH). Morphological alterations were noted in mice subjected to elevated doses of As_2_O_3_ (4 & 8 mg/kg bw). However, these changes were reversed in mice who received As_2_O_3_ + CUR co-treatment. Together, our findings provide preliminary evidence that curcumin protects mice striatal region from As_2_O_3_-induced behavioral, biochemical and morphological alterations.

## 1. Introduction

Arsenic (*As*), a naturally occurring metalloid, is ubiquitously distributed in the earth’s crust and ranks first in USEPA (United States Environmental Protection Agency) list of prioritized pollutants, thus posing a threat to human health. Arsenic poisoning is a matter of global concern exclusively in Latin America and Southeast Asian countries. Arsenic toxicity worldwide is primarily caused by consumption of contaminated drinking water or drinking water containing high levels of inorganic arsenic *(iAs)* [1–3]. Arsenic exists in both organic as well as inorganic forms with various oxidation states like elemental (0), trivalent (+3), pentavalent (+5) & Arsine (-3)[4–6].

Arsenic trioxide (As_2_O_3_), an inorganic compound of *As* has got notoriety in medical literature and is recognized as both poison and anti-cancer drug for the decades. Oncologists commonly use As_2_O_3_ as a first-line chemotherapeutic for acute promyelocytic leukemia (APL)[1,7]. The anti-neoplastic role of As_2_O_3_ in APL is due to As_2_O_3_-induced arrest of cell growth and apoptosis in NB4 cell line (APL cell line) or other cancerous cell lines [5,8,9]. As_2_O_3_ is the most active single agent for treating relapsed APL by inducing complete remission with few side effects. Currently, As_2_O_3_ is therapeutically being used in injectable form and commercially available under the trade name of Arsenox™ or Trisenox™.

Chronic exposure to inorganic arsenic has been associated with cognitive dysfunction in both rodents and humans. As it crosses the blood-brain barrier, it induces morphological alterations in the central nervous system (CNS) and may also result in differentiation syndrome when exposed environmentally or therapeutically [10,11]. As a result of its effective deposit in several brain regions, arsenic can cause various neurobehavioral disorders [12]. Arsenic is known to adversely affect the development of nervous system, cognitive functions, hearing, and may even result in peripheral neuropathy [13].

The accumulation of arsenic in CNS causes impaired neuronal activity, reduced neuronal migration, disrupted neuronal maturation and proliferation, and alteration of major neurotransmitter levels like dopamine, acetylcholine, serotonin, and glutamate[12,14,15]. Various studies have documented that arsenic-induced oxidative stress is one of the main mechanisms in *iAs*-induced neurotoxicity.

Basal ganglia are one of the brain regions lying deep inside the cerebral hemisphere, in close proximity to the internal capsule. Basal ganglia involve caudate & putamen nucleus, globus pallidus, substantia nigra, and subthalamic nucleus. Functionally, it controls cognitive tasks, locomotion, reinforcement-learning and goal-related task [16,17]. Any disturbance to the neurons of basal ganglia may result in difficulty in speech, balance & movements, and such symptoms are collectively referred to as “Parkinsonism”.

Clinical studies have shown the beneficial role of Antioxidant rich fruits & vegetables in reducing the incidence and intensity of various chronic diseases [18]. Commonly used Antioxidants are- Alpha Tocopherol, Melatonin, Vitamin A, Vitamin E, Taurine, N-Acetyl cysteine (NAC), Alpha lipoic acid and Curcumin [12,19]. Curcumin (CUR) is one of the major active herbal compounds which act as a potent antioxidant [20] and chelating agent.

Moreover, the therapeutic potential of curcumin supplementation has been reported in various neurodegenerative disorders [21,22], heavy metal toxicities [23], and oxidative tissue injuries. It acts as a chelating agent and potential scavenger of reactive oxygen species, thereby reducing oxidative stress (OS) in the tissues [24].

Numerous studies have demonstrated the Neuro-protective role of CUR in various neurodegenerative diseases like Alzheimer’s, Parkinson’s [25], and Cerebral ischemia [26]. Although there are numerous studies indicating the neurotoxic effects of arsenic and the role of CUR in attenuation of heavy metal neurotoxicity, however there is a paucity of in-vivo studies demonstrating the effects of varying doses of arsenic trioxide alone or in combination of curcumin on basal ganglia-mediated behavioural functions, oxidative damage, and neurotransmission in mouse striatal region.

Therefore, considering the reported role of As_2_O_3_ in the induction of oxidative stress (OS) and Neurodegeneration, present study attempted to determine whether antioxidant-curcumin could assist in mitigating neurotoxicity caused by As_2_O_3_ in the mouse striatum.

## 2. Materials and methods

### 2.1 Animals and treatment

Healthy adult male albino mice of Swiss strain weighing about 25 to 30g were used for the present study. All the drug treatments were carried out after obtaining ethical approval from the Institutional Animal Ethics Committee (IAEC). Guidelines of CPCSEA (Committee for the Purpose of Control and Supervision of Experiments on Animals) were strictly adopted for the use, care and handling of animals. The animals were initially acclimatized to the housing conditions for a week and then randomly segregated into various study groups as depicted in figure 1. The first group was control (Group I), which included the normal control group (Group Ia) that did not receive any treatment and the vehicle control group (Group Ib) received 5% aqueous solution of gum acacia (GA), the vehicle for Curcumin (CUR). The experimental groups comprised of Arsenic trioxide (As_2_O_3_) alone treated group (Group II), Curcumin (CUR) alone treated group (Group III) and As_2_O_3_ + CUR treated group (Group IV). Group II and IV were further divided into IIa, IIb, IIc & IVa IVb, IVc subgroups respectively based on different concentrations of As_2_O_3_ dosage i.e., 2, 4 and 8 mg/kg bw as shown in figure 1.

**Figure 1:**
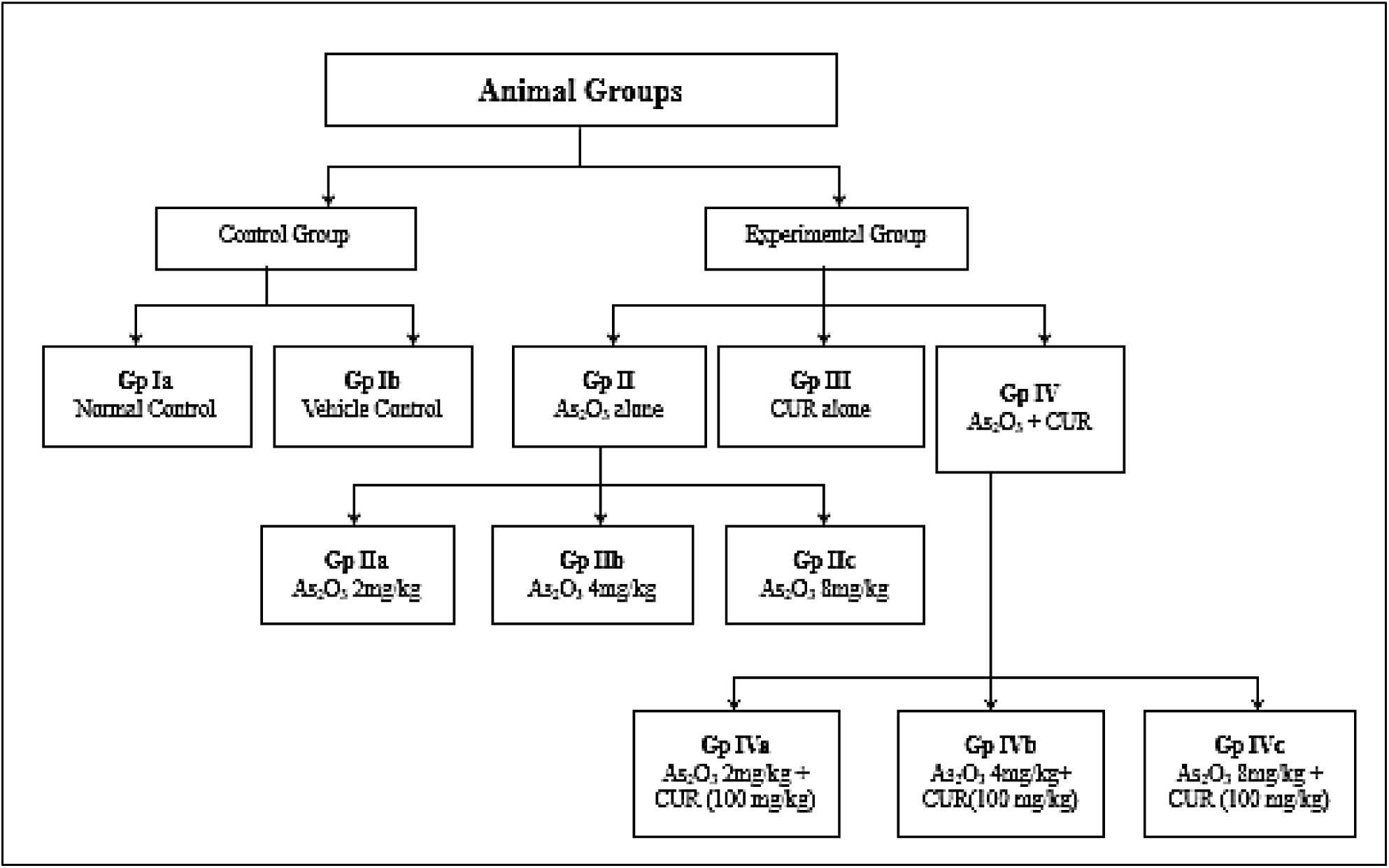
Flow chart illustrating various study groups and treatment regimen for As2O3 and Curcumin (Cur) for a period of 45 days

### 2.2 As_2_O_3_ and Curcumin dose regimen selection

Arsenic trioxide (As_2_O_3_), Curcumin (CUR) and Gum Acacia (GA) were procured from Sigma Chemicals (St. Louis, USA). The As_2_O_3_ treated groups alone received three doses of As_2_O_3_: 2mg, 4mg and 8mg/kg body weight on a daily basis all along the experimental period (45 days). The therapeutic dose (i.e., 2mg/kg bw) of As_2_O_3_ was calculated from the Human Equivalent Dose (HED) of 0.15 mg/kg bw, being routinely administered in APL (acute promyelocytic leukemia) patients over a period of 45 days (Iland & Seymour, 2013). The entire tested regimen (Arsenic trioxide, Curcumin and Gum Acacia) was administered through oral gavage for every animal group as mentioned above. Curcumin (100 mg/kg bw) was constantly administered to the curcumin alone group and curcumin co-administered groups (IVa, IVb & IVc).

### 2.3 Behavioral study

Over the course of 15 days of the experimental period, behavioral experiments were conducted to assess anxiety levels, general locomotor activity, motor coordination, memory & learning pertaining to basal ganglia as shown in figure 2A & B.

**Figure 2A:**
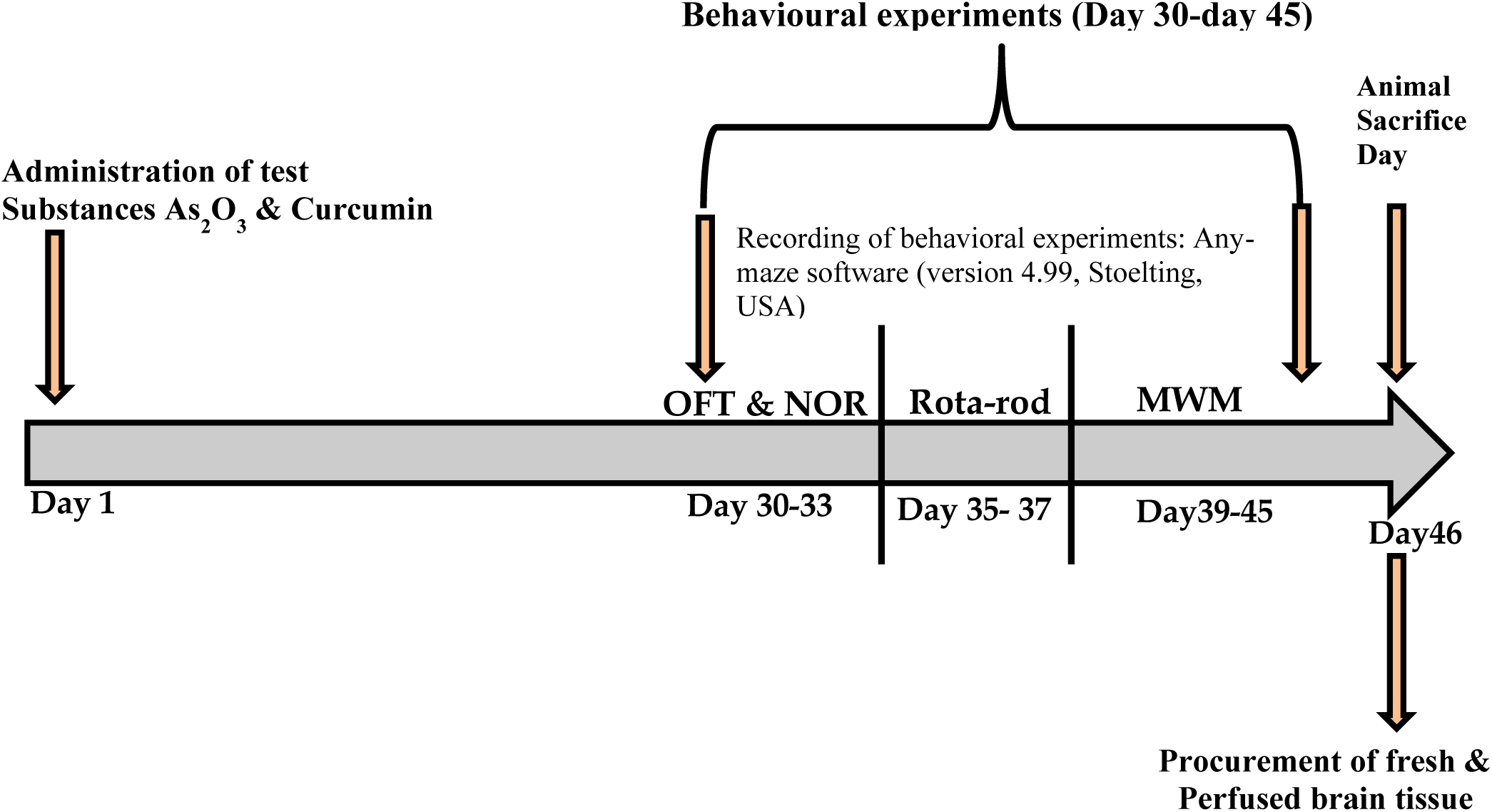
Timeline chart depicting the study progression.

**Figure 2B.**
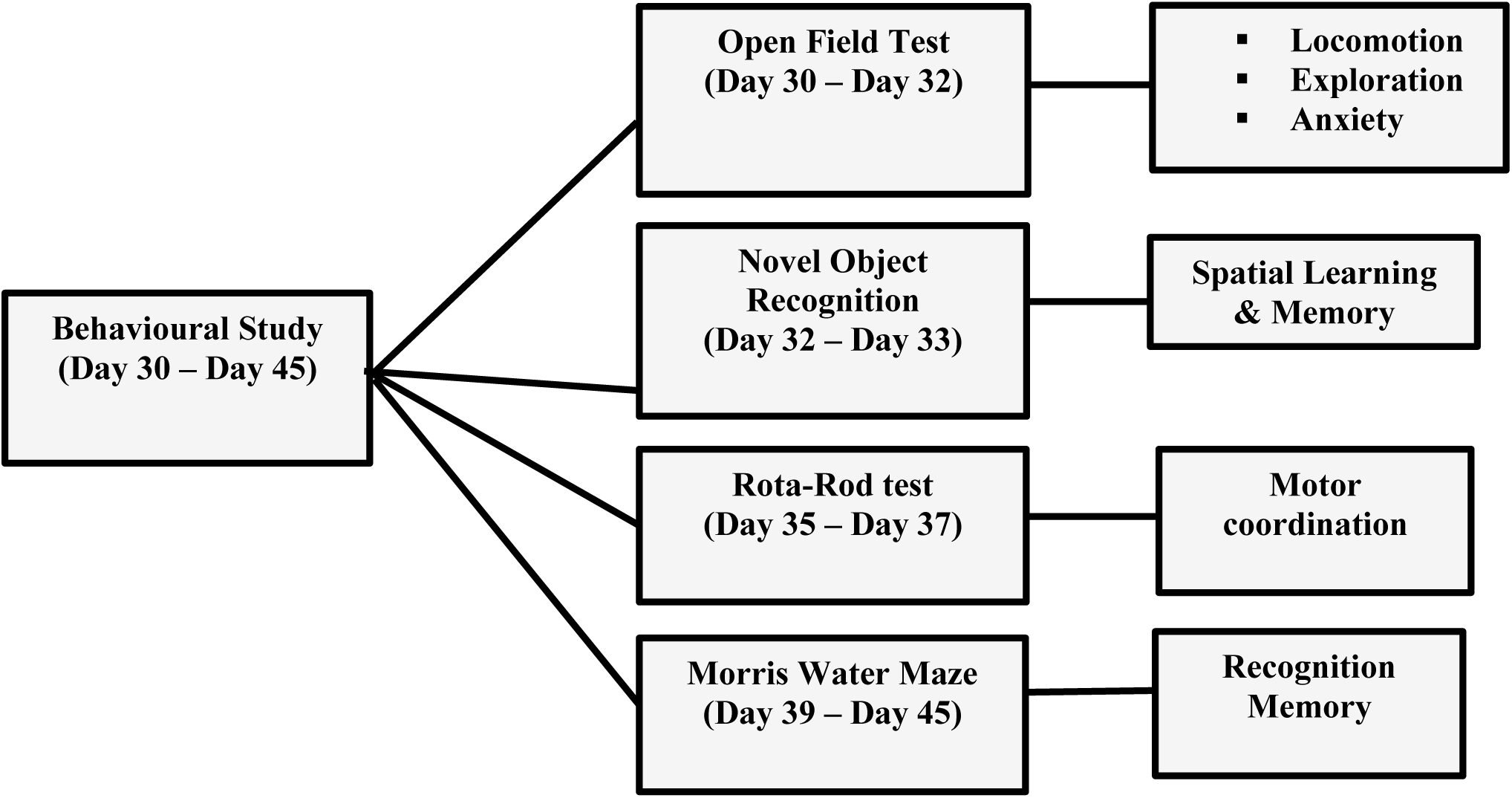
Work plan for behavioral study at the end of the experimental period (from day 30-day 45)

#### 2.3.1 Open field test (OFT)

OFT was performed from day 30 to 32 for evaluation of exploratory behavior & general locomotion activity & anxiety in rodents. OFT apparatus consists of a transparent plexiglass chamber with a large square arena which is marked virtually with grids and divided into four corners, peripheral and central zones. The test session began by placing the animals individually in the center of the arena and allowing them to freely explore the chamber for 5 minutes. The whole apparatus was connected to automated video tracking system with attached Any-maze software (Version 4.99). Meantime spent in the periphery & corner square of the chamber was recorded as a measure of anxiety i.e., more the time spent in the center of the field indicated lesser anxiety levels, on the contrary, more the time spent in the periphery & corner suggests higher anxiety levels, while total ambulatory time, distance travelled, velocity and no of line crossings were recorded as a measure of general locomotion & exploratory behavior of mice.

#### 2.3.2 Novel Object Recognition (NOR)

NOR was conducted from day 32 to 33 to study memory recognition in the mice. Briefly, the mice were first introduced to two similar objects during the habituation phase (training). On the test trial day, one of these objects was substituted with a novel object that differs in size, shape and color. The measure of the test is the amount of time spent by rodents exploring each object. It is based on novelty preference test and calculated in the form of recognition index (RI) or discrimination index (DI) [27].

#### 2.3.3 Rotarod test

(day 35 to 37) was performed to assess motor coordination and balance of the rodents. In this experiment the mice were allowed to maintain balance on an accelerating rod at a speed ranging between 4 to 40 rpm for 300 seconds. The test comprised four trials (T) performed across two consecutive days (Day 1: T1 and T2; Day 2: T3 and T4). The latency to fall for each mouse was recorded manually i.e., time point at which the mouse falls off from the rotating rod.

#### 2.3.4 Morris water maze (MWM)

MWM test is widely used for the assessment of spatial learning and memory-like abilities in rodents. This test was performed for 7 consecutive days (days 39 to 45), consisting of habituation (Training/visible platform) phase, acquisition phase (hidden platform) and retention phase (Probe trial/without platform). Apparatus consists of a circular pool, divided into four equal quadrants (NE, SE, SW, NW). A platform was placed in one of the target quadrants (SW). The test is based on the principle that rodents can learn to swim from any starting point towards the hidden/visible platform using external distal cues, reflecting learning and memory. The test session began by releasing the animal gently into the pool at water level from diverse start locations (N, E, SE, NW) facing the tank wall. Each animal received 4 trials per day and were allowed to explore the water-maze and locate the submerged platform in each interval of 60 sec. Escape latency (seconds) was recorded by the ability of the animal to locate submerged platforms in the water maze. If it surpasses 60 sec, then, it is recorded as 60 sec, and the animal is manually guided to the platform with the help of a wooden stick. The parameters recorded during acquisition phase and retention phase (Probe trial) included escape latency (the time taken by the animal to locate the submerged platform from its start location), mean Distance traveled (m), mean speed (m/sec), time spent in target quadrant (where platform was positioned) and number of platform crossings during probe trial [28].

### 2.4 Animal sacrifice and tissue procurement

Following behavioral assessments, the animals were sacrificed on day 46, either by perfusion fixation to get fixed tissue or by euthanasia to get fresh tissue. For perfusion fixation process, 4% of the paraformaldehyde was prepared in 0.1M phosphate buffer (PB) and administered transcardially after (2–3 min.) anesthetizing the animal with diethyl ether [6,29–32]. The brain specimens were carefully removed and then immersed in the same fixative for post fixation. The perfusion fixed brains were processed for paraplast embedding and cryo sectioning. CV staining was performed on paraplast sections for studying the morphological and morphometric features. While fresh basal ganglia tissue were procured for the biochemical estimation after euthanizing the animals.

For obtaining fresh brain tissue, the mice were subjected to cervical dislocation [33,34] and the brains were dissected out immediately, blanched in ice-cold isotonic saline (0.9%) and then weighed. The brain was placed on the ice pack and the striatum region dissected out from the brain. This striatum tissue was processed for estimation of antioxidant markers (GSH & GPx) & oxidative stress markers (MDA) and Western Blot analysis. For biochemical estimation, striatum tissue (40 mg) were sonicate in 0.1M PB (pH =7.4) (VCX 500, Sonics-USA) for 1 minute at 4°C to obtain 10% w/v homogenate [35,36]. The prepared homogenate was then centrifuged for 15 minutes @ 10,000 rpm at 4°C and then the supernatant aliquots were prepared and stored at -80°C till further use. The level of antioxidant markers glutathione (GSH) and Glutathione peroxidase (GPx) and the end product of Lipid peroxidation process (MDA) were determined in the supernatant following specific protocols.

### 2.5 Biochemical Estimation

#### 2.5.1 Determination of Reduced Glutathione (GSH)

GSH levels were measured the method of Ellman (1959) which is based on the optimized enzymatic recycling employing the Ellman’s reagent (DTNB) and the glutathione reductase enzyme. Glutathione reductase enzyme decreases the level of GSSG to GSH while the DTNB (5-5’-dithiobis [2-nitrobenzoic acid]) reacts with GSH to form GS-TNB and TNB and is yellow color chromophore with maximum absorbance at 415 nm. Then the glutathione reductase reduces the GS-TNB to GSH and TNB by, thus this enzymatic recycling of GSH improves the sensitivity of the experiment.

The tissue supernatants were initially de-proteinized by equal volume of freshly prepared 5% TCA in striatum homogenates (100µl). This mixture was vortex and centrifuged at 5000 rpm for 5 min at 4°C and the clear supernatant (sample) was obtained [38,39]. The standards for reduced glutathione were prepared in series of known dilutions (1-2000 µg/ml) using 0.3M PB (pH=8). PB was also used for blank measurement. 30µl of supernatant (sample) was pipetted into wells A-H (column 5,6) of 96 well-plate. Equal amounts of 0.3M PB were added in wells A of column 1,2 and equal volume of standards were pipetted into wells B-H (columns 1,2) and A-E (columns 3,4) of 96 well-plate. Then 50µl of 0.3M phosphate buffer was added in both samples as well as standards followed by 150µl of DTNB (5,5’-dithiobis 2-nitrobenzoic acid in 1% trisodium citrate known as Ellman’s reagent). Within 10-15 min, the absorbance (OD) of the samples was measured at 412 nm by microplate reader (Synergy H1, Biotek). A standard curve was drawn by plotting known concentrations of standards along X-axis against their absorbance along Y-axis. The linear equation [x=(y-b)/m] was used to calculate the unknown protein concentration in the samples where y = signal (OD); m = slope; x = protein concentration; b = (y intercept) constant [40]. GSH concentration in the samples was expressed as µg/g of wet tissue. All the measurements were performed in duplicate.

#### 2.5.2 Determination of Glutathione peroxidase activity (GPx)

GPx levels in brain were determined by the method developed by Flohe and colleagues [42]. Briefly, striatal tissue was homogenized in 0.1M Pb (pH 7.4) followed by centrifugation at 1500g for 10 min at 4°C. Then, the supernatant was collected in another MCT (micro-centrifuge tube) and again centrifuged (10,000g) at 4°C for 30 min. The double centrifuged supernatant was obtained and used for the assay of GPx activity. The final volume of the reaction mixture obtained was 1 ml which constituted 0.1M Pb (pH 7.4), 2mM GSH, sodium azide (10 mM), hydrogen peroxide (1 mM) and then incubated at 37°C for the period of 15 min. TCA (10%) was added to stop the reaction. The reaction mixture was centrifuged for 5 min at 1,500×g to settle down the protein at bottom and the supernatant was transferred to another tube having 0.2 ml of 0.1M Pb (pH 7.4) and 0.7 ml of DTNB (0.4 mg/ml). The absorbance was read at 420 nm after vortex the tubes. The results were expressed as nmol of GSH oxidized per minute per mg of protein.

#### 2.5.3 Determination of lipid peroxidation (MDA)

To measure the end product of lipid peroxidation, malondialdehyde (MDA) estimation was carried out in the caudate putamen region [43]. This experiment employs the reaction of a chromogenic reagent with MDA. For this procedure the tissues were homogenized in 0.1M Pb (pH 7.4). The homogenate was incubated with SDS (10 %, w/v), TBA (0.8%) and 20% acetic acid for one hour in boiling water bath. The intensity of pink color formed by the chromogenic reagent and recorded at 532 nm. The total amount of TBARS was calculated using a molar extinction coefficient of 1.56 x 10^5^ M cm^-1^. The results were expressed as the amount of MDA (µmol) formed per hour per mg of protein.

### 2.6 Morphological observations

The perfusion fixed brain tissues were processed for paraffin embedding following the conventional protocol. Briefly, the tissues were washed overnight under running tap water. Following day, tissues were dehydrated in ascending grades of alcohol [70% (1 hour), 80% (40 min.), 90% (20min.), 2 changes of 96% (15 min. each)] Following dehydration, the brain specimens were cleared in cedar wood oil until the tissue becomes transparent. For adequate infiltration, the brain specimens were subjected to three changes of paraplast for 30 minutes each at 56°-58°C in incubator. The embedded tissue blocks were sectioned coronal at 7 µm thickness on rotary microtome (Shandon AS325). The sections were later subjected to Cresyl Violet (CV)/nissl staining and observed under bright-field microscope.

Thorough morphological study and morphometric analysis was carried out in CV stained paraplast sections of AOI i.e., Striatum (Caudate putamen) of control and experimental animals. The Striatum (Figure 3 A & B) was identified from the anatomical landmarks in coronal sections of brain as presented in the Stereotaxic Mouse Brain Atlas (Paxinos and Franklin, 2008). Morphological observations were carried out on the CV-stained coronal serial sections (7μm thick) of caudate putamen (Bregma +0.98mm to 0.02mm) [44] region using bright field Nikon E-600 microscope equipped with Nikon Digital Camera System (DS-Fil-U2) as shown in figure 4. The CV-stained coronal sections from all groups were observed under higher magnification (40X) and various morphological features like shape and size of the neurons, shape and location of the nuclei and the number of nucleoli per nucleus were considered. Cell density was determined both qualitatively and quantitatively. Every 8^th^, CV-stained coronal serial sections (7μm thick) of caudate putamen region were taken. Whole region of Caudate Putamen were divided into DM, DL, VM & VL to measure the area of neurons and the centre of area is selected in each section and Grid frame 157*157 were formed to measure the neuron size [45]. Only those neurons were selected for measurement, which were had distinct nucleolus and two to three nucleoli [46].

**Figure 3:**
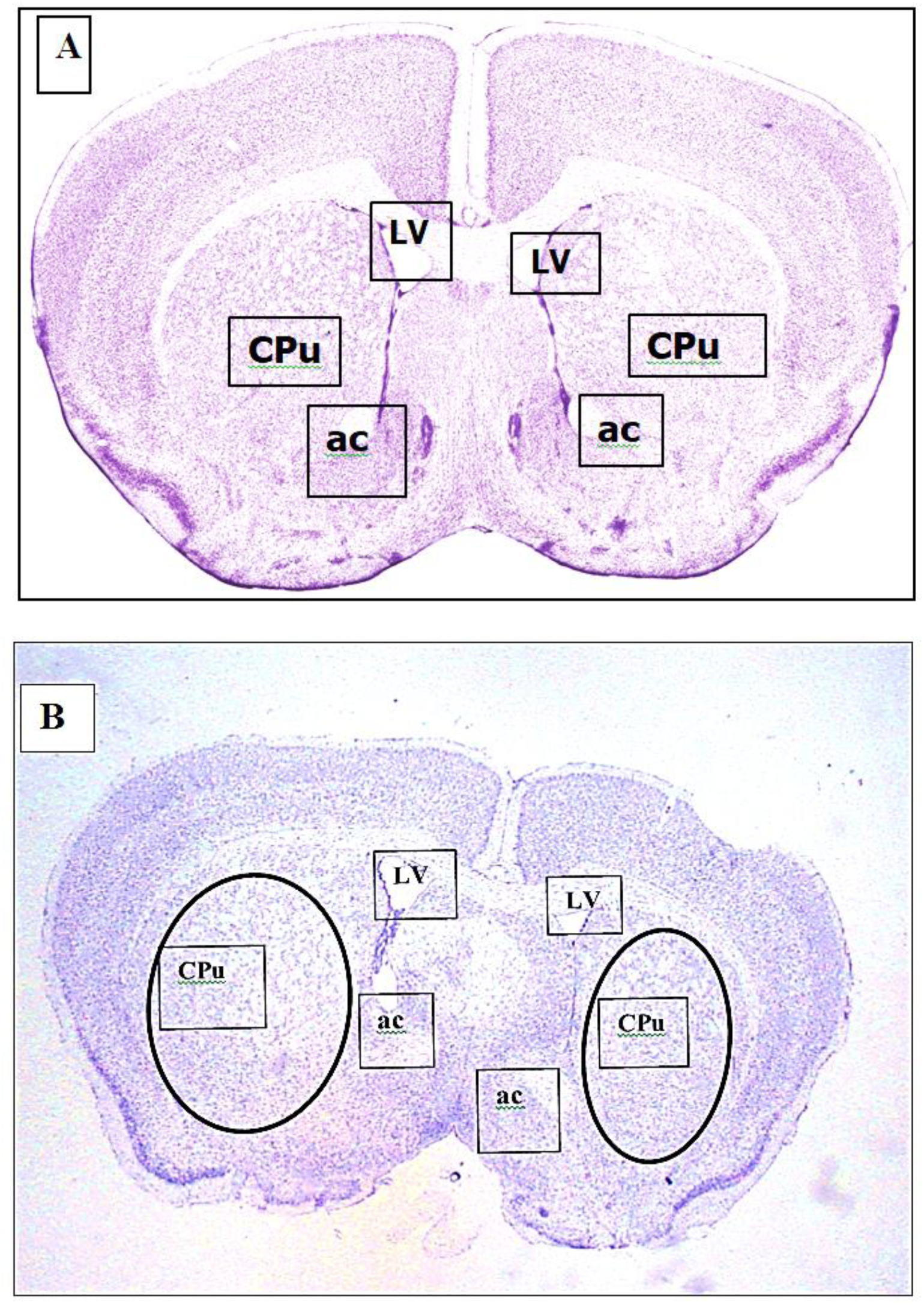
A. Photomicrograph showing reference image adopted from Mouse Brain Atlas by Keith BJ Franklin & George Paxinos (Bregma level: +0.98 mm). B. Photomicrograph showing CV-stained coronal section of mouse caudate putamen (Cpu) region and the anatomical landmarks (Bregma level: +0.98 mm). LV-Lateral ventricle; ac-Anterior commissure.

**Figure 4:**
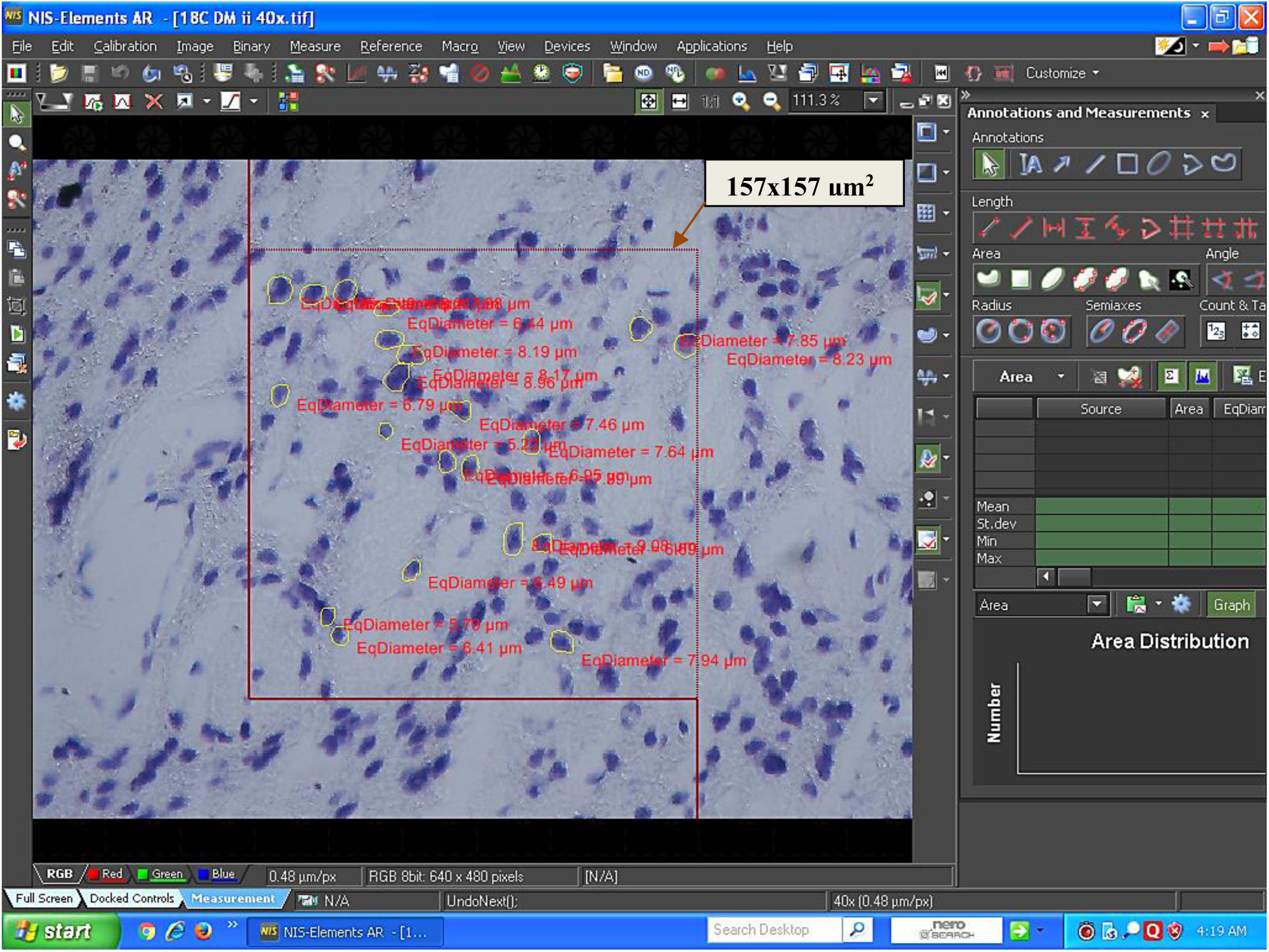
Snapshot of screen showing rectangular frame (157×157 μm^2^) superimposed on one of the reference areas in caudate putamen in NIS Element AR3.10 image analysis software.

### 2.7 Statistical analysis

The data was compared among various study groups (Control, As_2_O_3_ alone and As_2_O_3 +_ Curcumin co-treated) using appropriate statistical tests. The data was analyzed, and graphs were plotted by using Stata 15.1 (Stata Corp LLC, USA) and GraphPad Prism 8 (V.8.4.3, San Diego, California). Data was represented as mean ± SEM/SD or median (where applicable). The normality of the data was checked using the Shapiro Wilkis test. For normally distributed data, one-way ANOVA was applied followed by Bonferroni test to compare the differences among various groups. The Kruskal-Wallis test followed by pairwise comparison (Bonferroni method) was applied to compare the medians of behavioral tests among the groups for non-normal data. Escape latency, mean speed and distance traveled in MWM were compared using repeated measures ANOVA among the groups over a period since the data was correlated. P-value ≤0.05 was considered statistically significant.

## 3. Results

### 3.1 Effects of CUR on general appearance of mice exposed to As_2_O_3_ treatment

All the animals were observed routinely for their body weight and physical signs such as state of alertness, grooming, hair loss and any injury during the experimental period (day 1 to 45). No obvious alterations were seen in the general appearance (hair loss, any external injury) and physical activities (gate pattern, rearing, grooming) among the control and the experimental groups receiving low (2 mg) and medium (4 mg) doses of As_2_O_3_. However, in animals receiving higher doses of As_2_O_3_ (8mg/kg bw), patchy hair loss was observed on dorsal and ventral surfaces of the body. The values for the percentage body weight of the control and the experimental animals on day 1 & 45 are shown in supplementary table 1(S1). On day 1 of the experimental period, the mean values of the body weight of the control and the experimental animals were almost similar. Following the two weeks of the experimental period, a gradual increase was observed in the body weight of almost all the animal groups. Moreover, at the end of the experimental period, no significant difference (p<0.05) was observed between the control and experimental groups.

### 3.2 Effects of As_2_O_3_ +CUR on basal ganglia-mediated behavioral functions on exposed mice

The behavioral experiments were performed to evaluate the effects of As_2_O_3_ exposure at different concentrations (2mg, 4mg & 8 mg/kg bw) or along with curcumin on basal ganglia-mediated behavioral functions. A battery of behavioral tests was conducted from day 30 to day 45 of the experimental period. The recording of all the behavioral experiments were carried out by using Any-maze software (version 4.99, Stoelting, USA).

#### 3.2.1 Open Field Test (OFT)

The median velocity in the open field arena was found to be comparable across the control and experimental groups. It was observed that the median values of velocities in groups treated with As_2_O_3_ alone were lower than the control groups whereas co-administration of CUR resulted in comparative increase in velocities which shows the improvement of locomotor activities as compared to the corresponding As_2_O_3_ alone treated groups (Fig 5A & S2).

**Figure 5:**
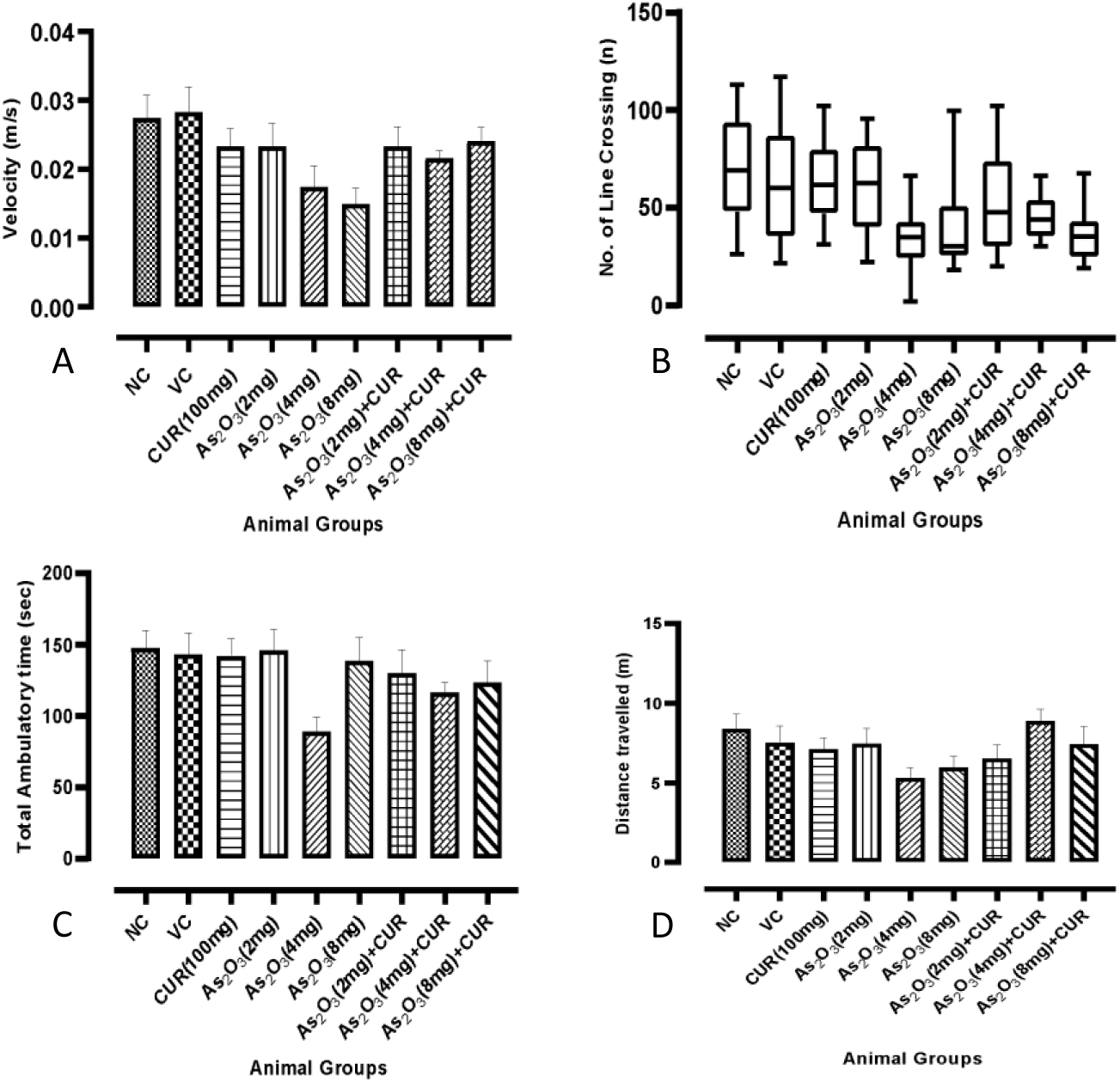
Open Field Test (OFT) (n=12/group) across control and experimental groups. A. velocity (m/s), B. Number of line crossings, C. Total ambulatory time (sec), D. Total distance travelled (m)

The numbers of line crossings in the open field were found to be comparable among the animal groups (Fig 5B & S3). Apparently, the numbers of line crossings by As_2_O_3_ alone treated were lesser as compared to normal and vehicle control groups but the difference was not statistically significant. The co-administration of CUR resulted in an increase in number of line crossings as compared to the corresponding As_2_O_3_ alone treated groups.

The mean values for number of line crossings, ambulatory time and total distance traveled were found to be lesser in experimental groups as compared to the control groups (Fig 5B-D & S4, S5). This decrease in locomotors’ activity was found to be more prominent in groups treated with higher concentration of As_2_O_3_ (8mg) which was considerably improved after co-administration of CUR.

When compared across the groups the time spent in the center area was significantly decreased in As_2_O_3_ 8mg alone treated group as compared to control group (P<.0016), a significant association was found. Co-administration of CUR considerably increased the median time spent in Centre in case of 4mg and 8mg As_2_O_3_ alone treated groups (Fig 6A & S6).

**Figure 6:**
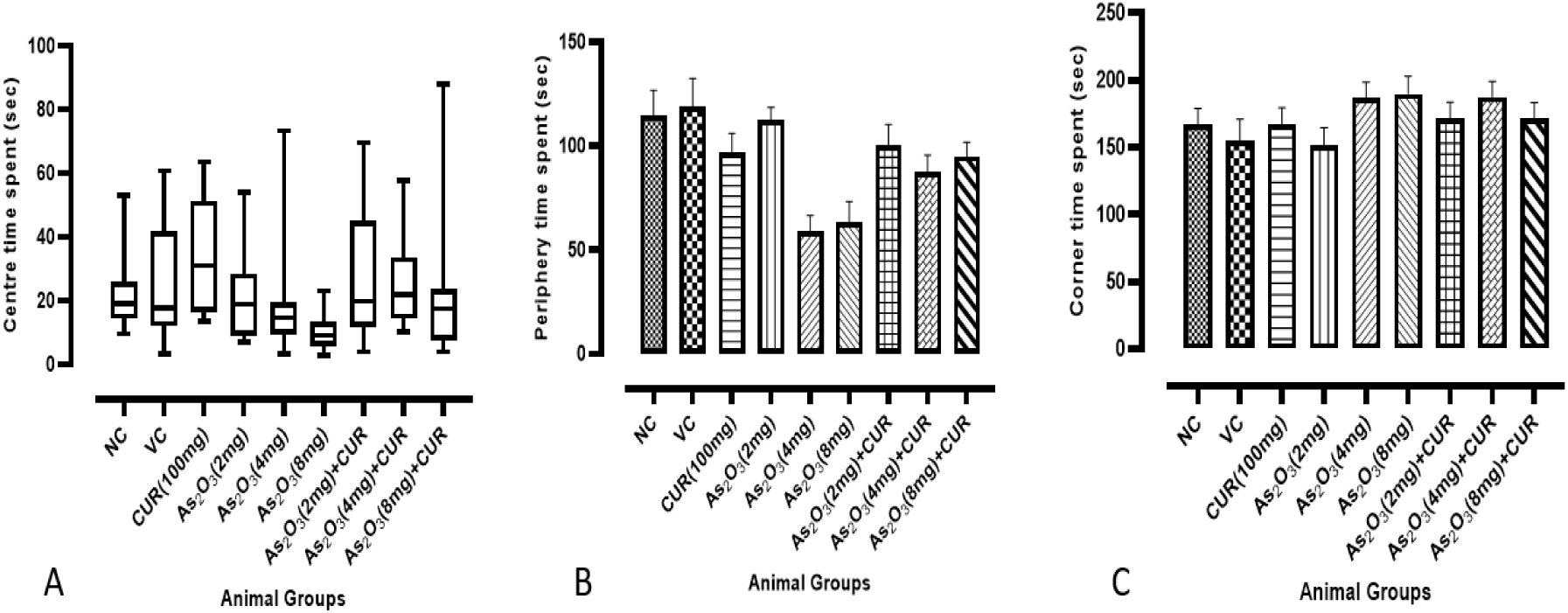
The diagrams showing the time (sec) spent in center (A) and Bar diagrams showing the periphery (B) and corner (C) time spent in the open field by control and experimental groups.

The mean time spent in the peripheral zone of open field arena ranged from 59.01±26.43-118.82±47.4 second and was also comparable across the groups (P<.0001). When compared across the groups the time spent in peripheral zone was also significantly decreased in As_2_O_3_ 8mg alone treated group as compared to control group (P<.003), Co-administration of 4mg and 8mg As_2_O_3_+CUR treated groups with the mean value 87.49±28.35 & 95.04±23.48 second respectively found to be considerably increased the mean time spent in peripheral zone. As depicted in Fig 6B &S6.

When inter-group analysis was done the time spent in corner zone was observed to be decreased in As_2_O_3_ 2mg alone treated group with the mean value 151.4±45.63 as compared to control group, whereas co-administration of As_2_O_3_ (2mg)+CUR treated group showed increased time spent in corner with the mean value of 171.41±40.93 second found as compare to As_2_O_3_ 2mg alone treated group (Fig 6C & S6).

The center entries were found to be comparable across all the groups (p<.0013). We have observed that the 8mg As_2_O_3_ alone exposed group with median value 3(1-4.5) showed significantly decreased center entries in open field test (p<.0034) as compared with control groups as shown in Fig 7A & S7. Similarly, 4mg As_2_O_3_ alone treated groups with median value 4.25(1-13) showed decreased center entries as compared with control groups.

**Figure 7:**
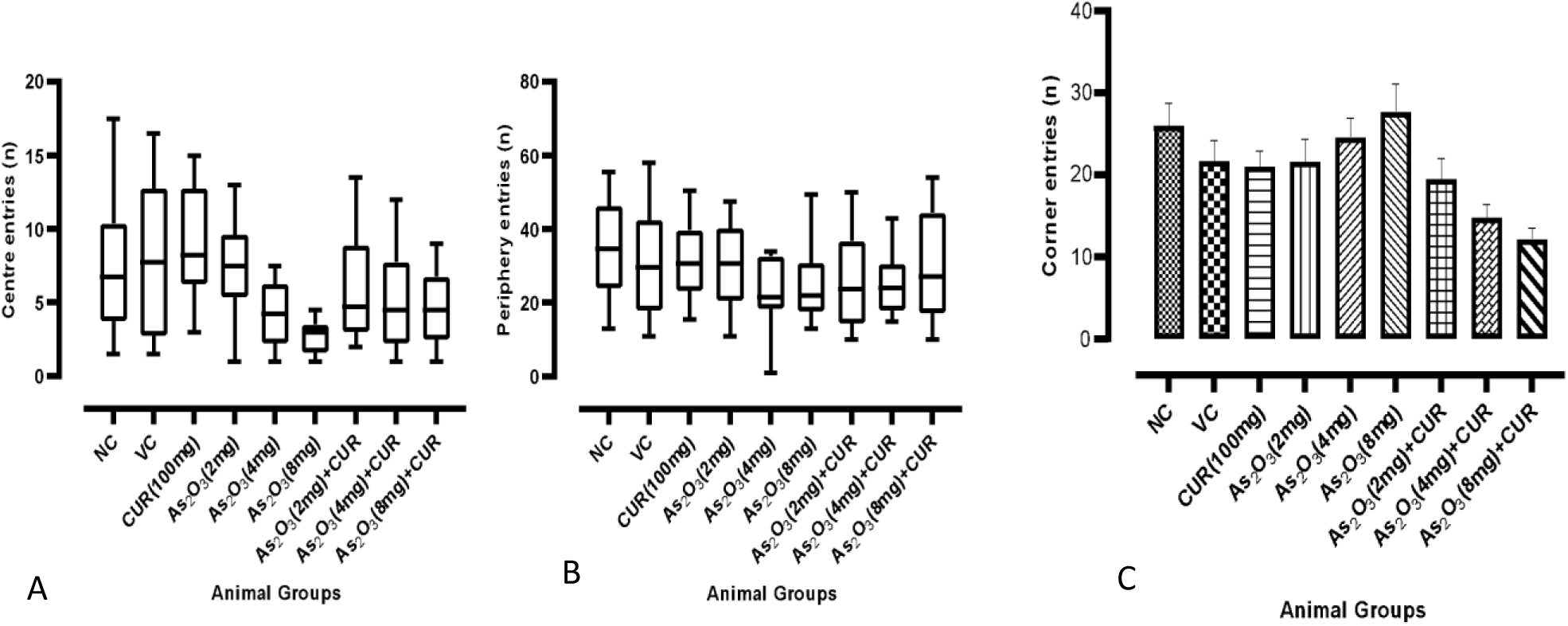
The number of center entries (A) periphery entries (B) and the corner entries (C) in control and experimental groups during OFT (n=12 animals/groups).

The median entries in the peripheral zone of the open field arena ranged from and were also comparable across the groups (p<.490). The median value of time spent in the periphery is depicted in Figure 7B & S7. When compared across the groups the entries in the peripheral zone were decreased in As_2_O_3_ 2mg, 4mg & 8mg alone treated group as compared to control group. Whereas co-administration of 4mg and 8mg As_2_O_3_+CUR treated groups with the median values of peripheral entries 24(15-43) & 27.25(10-54) respectively were found to be considerably increased, however failed to reach the statistical significance.

When pairwise comparison was done across the groups, the entries in corner zone was observed to be significantly decreased after co-administration of 8mg As_2_O_3_ with CUR showing the mean values of 12.16±4.46 in 8mg As_2_O_3_ alone exposed group with the mean value 27.7±11.75 (P<.0001) as shown in Fig 7C & S7. Similarly, it was observed in co-administration of 2mg & 4mg As_2_O_3_+CUR treated group with the mean value of 19.5±8.59, 14.79±5.51 respectively as compared to 2mg & 4mg As_2_O_3_ alone exposed groups with mean values of 21.62±9.37 & 24.62±8.02 respectively.

#### 3.2.2 Novel Object Recognition (NOR) Test

To address the changes in recognition memory following As_2_O_3_+CUR co-treatment, we conducted Novel object recognition test in mice. In NOR task, we recorded recognition index (RI) & discrimination index (DI) as the major indices of recognition memory. Animals tend to have an innate preference for novelty such that if they recognize the familiar object, then they will spend most of their time exploring the novel object over the familiar object.

The median RI in NOR test ranged from 0.33(0-0.92) to 0.66(0-1) and was also comparable across the groups (P<0.2595). The median value of memory RI in median (min-max) is shown in Figure 8A & S8. When compared across the groups the memory recognition index was observed to be decreased in 2mg, 4mg & 8mg As_2_O_3_ alone treated groups with the median values 0.59(0-0.95), 0.38(0-0.75), 0.33(0-0.92) respectively as compared to control group with the median value 0.66(0-1). Whereas co-administration of 2mg, 4mg & 8mg of As_2_O_3_+CUR treated groups with the median values of 0.5(0.32-0.92), 0.56(0-1), & 0.36(0-1) showed increase in memory RI as compared to 2mg, 4mg & 8mg As_2_O_3_ alone treated groups with the median values 0.59(0-0.95), 0.38(0-0.75), 0.33(0-0.92) respectively.

**Figure 8:**
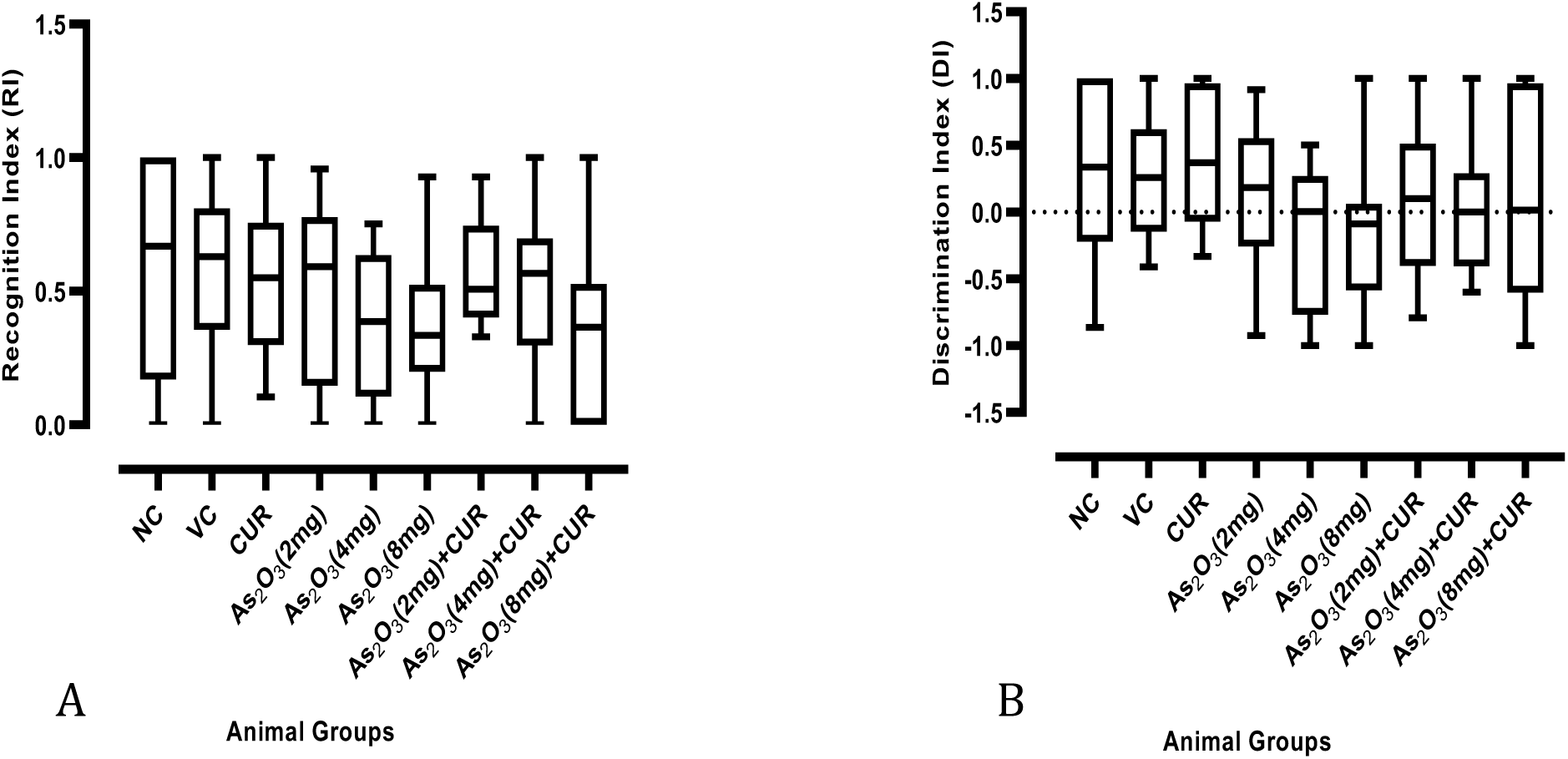
The recognition (A) and discrimination (B) indices across control and experimental groups (n=12 animals/groups) in NOR test.

The median DI in NOR test ranged from -0.08(-1-1) to 0.37(-0.33-1) and was also comparable across the groups (p<0.2696). The median value of object DI in median (min-max) shown in Figure 8B & S8. When compared across the groups the object DI was observed to be decreased in 2mg, 4mg & 8mg As_2_O_3_ alone treated groups with the median value 0.18(-0.92-0.91), 0(-1-0.5), -0.08(-1-1) respectively as compared to control group with the median value 0.33(-0.86-1). Whereas co-administration of 2mg, 4mg & 8mg of As_2_O_3_+CUR treated groups with the median value of 0.1(-0.79-1), 0(-0.6-1), & 0.01(-1-1) showed increase object discrimination as compared to 2mg, 4mg & 8mg As_2_O_3_ alone treated groups with the median value 0.18(-0.92-0.91), 0(-1-0.5), -0.08(-1-1) respectively.

#### 3.2.3 Rota-Rod Test

Rota-rod test is widely used to evaluate motor co-ordination, balance, and motor learning in rodents. The test included-time spent by animal in balancing itself on the accelerating rod before falling. The median latency to fall in Rota-rod experiments ranged from minimum 10.25(2.37-37.2) second to maximum 36.88(14.9-69.37) second, also comparable across the groups (p<.0001) and the values are shown in Fig 9 & S9. In our observation we have found that 2mg, 4mg & 8mg As_2_O_3_ alone treated group with the median value 17.01(2.5-39.6), 10.25(2.37-37.2) & 11.76(2.27-37.97) seconds respectively showed significant reduction in latency to fall, when compare with control groups with the median value 36.23(20.2-64.33) sec. Whereas co-administration of 2mg, 4mg and 8mg As_2_O_3_+CUR treated groups with the median value 21.65(10.9-66.3), 24.81(12.5-38.47) & 19(11.27-37.2) seconds respectively were found to be considerably increased latency to fall when compare with 2mg, 4mg & 8mg As_2_O_3_ alone treated group with the median value 17.01(2.5-39.6), 10.25(2.37-37.2) & 11.76(2.27-37.97) seconds respectively but does not reach upto significant level (p>0.008).

**Figure 9:**
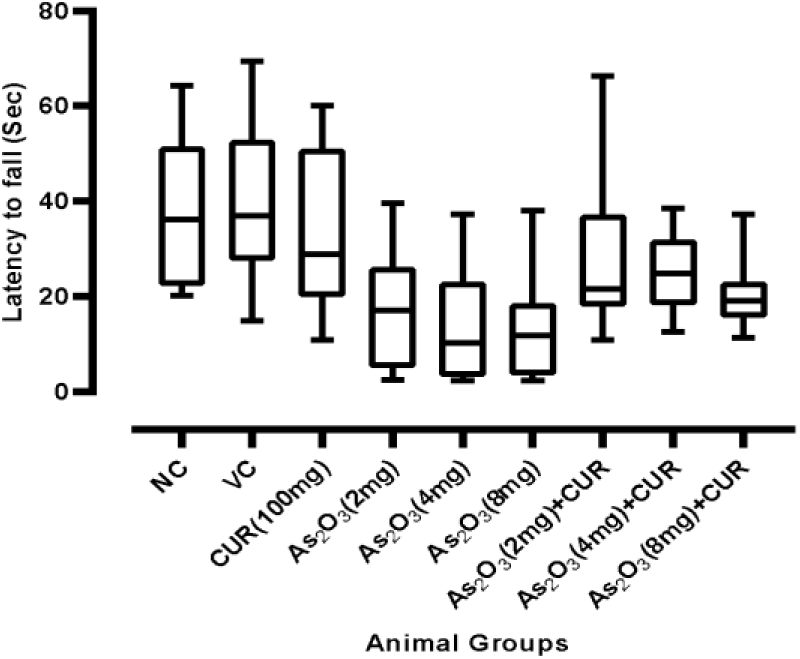
The latency to fall (sec) in various control and experimental groups (n=12 animals/group) in Rota rod experiment.

#### 3.2.4 Morris Water Maze Test

Morris water maze test (MWM) was used for the assessment of spatial learning & memory abilities in mice. The test was divided into three phases including habituation phase (Days 1 & 2), Acquisition phase (days 3-6) and probe trial (Day 7). The average escape latency, distance travelled, and swim speed were recorded during acquisition phase (hidden platform), whereas time spent in the platform quadrant (SW) and number of platform crossings (n) were recorded on the day of probe trial (without platform).

### Spatial acquisition phase

The learning ability of animals was evaluated from escape latency (EL-time taken by mice to find the hidden platform), path length (distance travelled to find the platform) and average swimming speed (locomotor activity). The average escape latency was observed to be gradually decreasing in successive trials in all the groups but found to be highly significant among control and experimental groups (p<.0006).

On day 1 of acquisition phase, the controls, As_2_O_3_ alone treated and As_2_O_3_ + CUR co-treated mice spent almost similar time in locating the hidden platform. The EL values on day 4 in the As_2_O_3_ + CUR co-treated groups (2, 4 & 8 mg/kg + 100 mg/kg) showed a significant (p<.0006) decrease (fig. 10 & S10). Overall, the animals administered CUR along with As_2_O_3_ reached the platform in apparently shorter time on day 4 of acquisition phase as compared to the As_2_O_3_ alone exposed groups.

**Figure 10:**
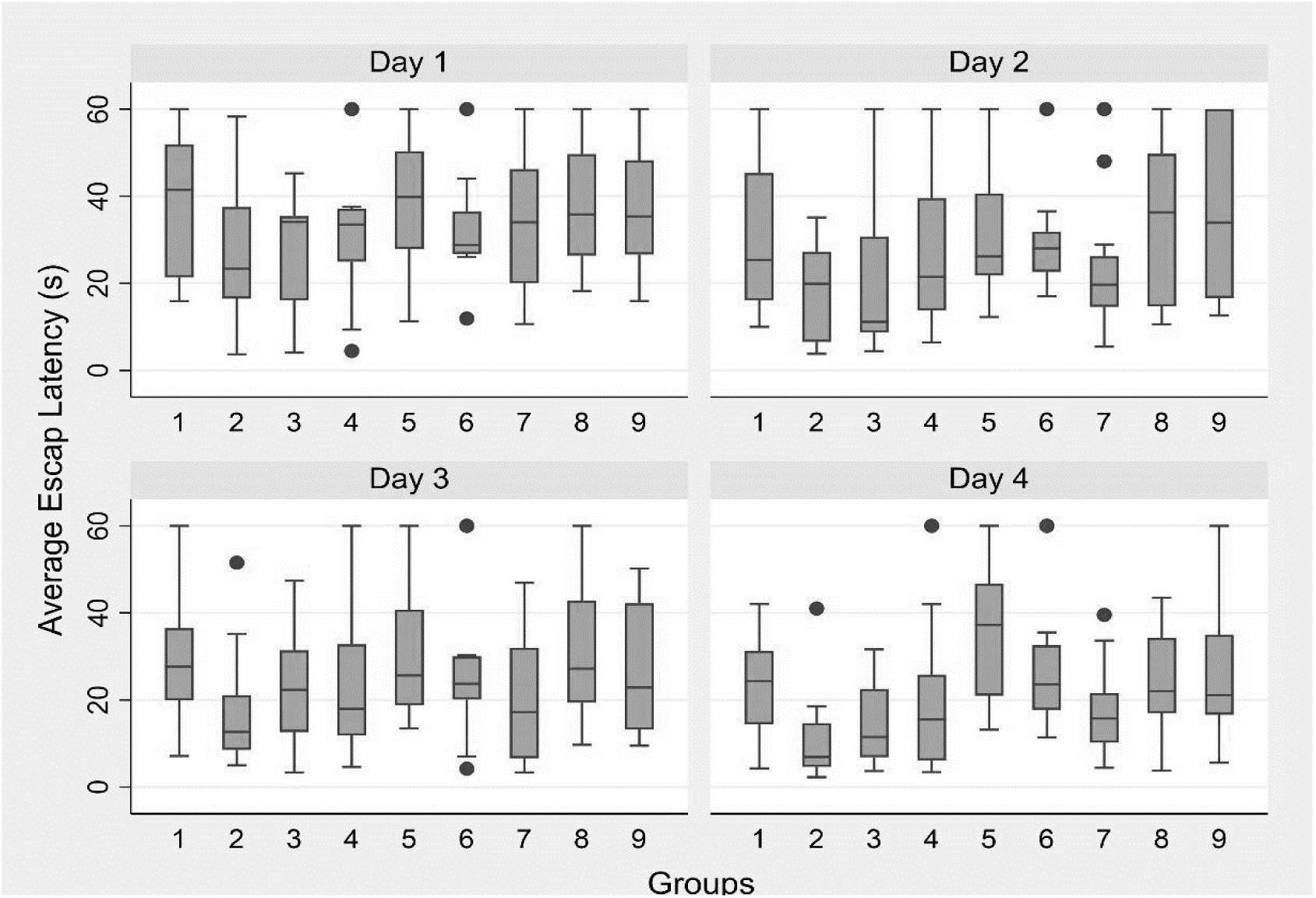
Average escape latency (n=12/group) in MWM during acquisition phase. Numbers 1 to 9 along x-axis denote the study groups.

Similarly, the distance travelled to reach onto the hidden platform by the control and experimental animals showed no differences (p<.5511) on day1 of acquisition phase. While on subsequent days (days 2 &3) of acquisition phase, animals receiving As_2_O_3_ alone (2, 4 & 8 mg/kg) travelled longer distance as compared to the shorter distances travelled by controls and curcumin supplemented groups (2, 4 & 8 mg/kg + 100 mg/kg) but the difference was not significant among the groups (p<.0174, p<.0856 respectively), but on 4^th^ day of acquisition period the distance travelled by control curcumin co-administered groups significantly more as compared to As_2_O_3_ alone (2, 4 & 8 mg/kg) treated mice (p<.0017) as shown in Figure 11 & S11. The As_2_O_3_ alone (2, 4 & 8 mg/kg) treated mice travelled a shorter distance on day 4 to reach the platform as compared to day 1 but the difference was not statistically significant. These results do suggest CUR supplementation induced substantial improvement in spatial learning and memory abilities over the four days of acquisition phase in As_2_O_3_ + CUR co-treated groups. In the similar pattern, no significant difference was observed in swimming speed among experimental groups over a period of four days; thereby, indicating no evidence of motor disturbance or alterations in motivational and visual abilities in various groups (Fig 12 & S12).

**Figure 11:**
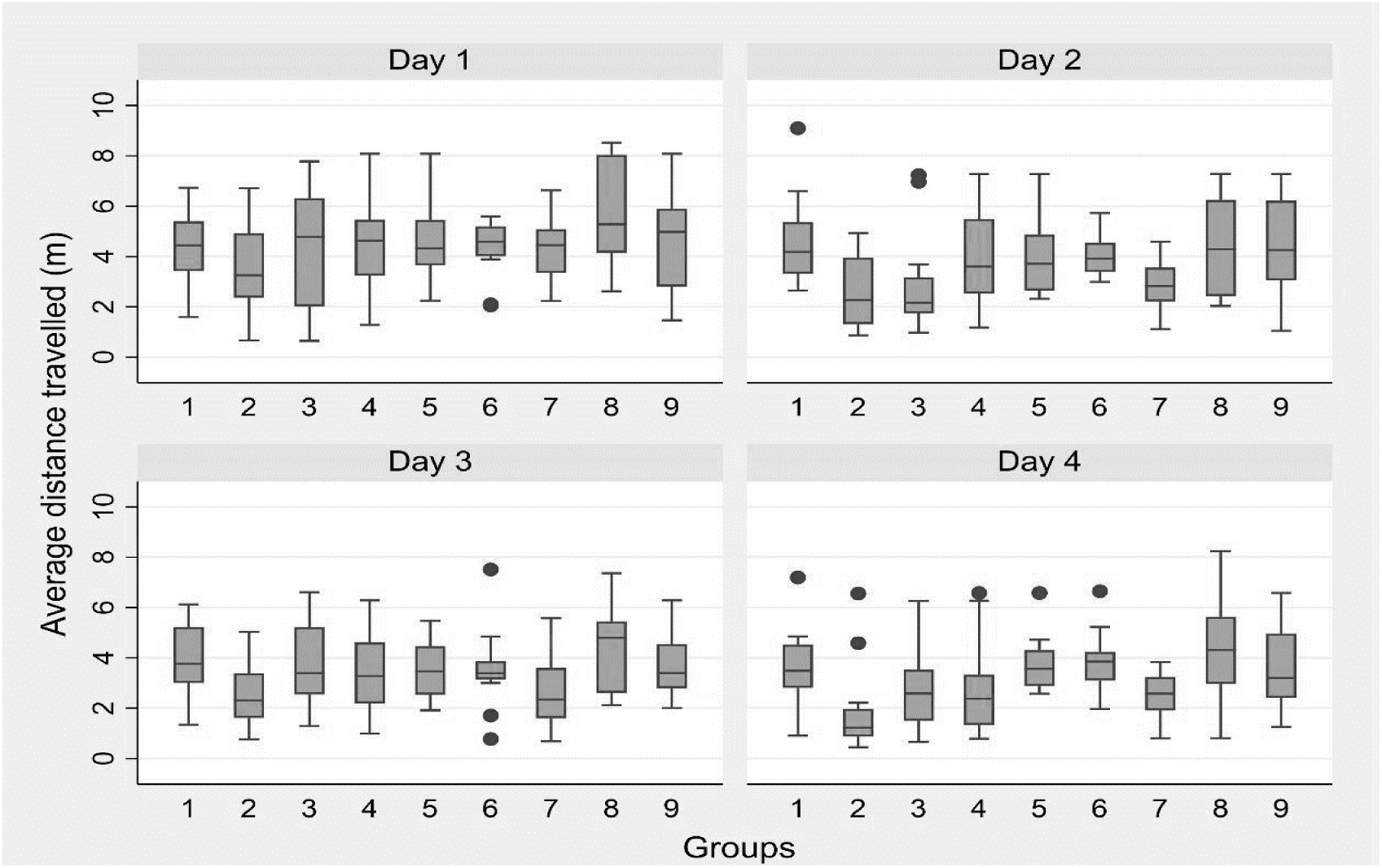
Average distance travelled (n=12/group) in MWM during acquisition phase. Numbers 1 to 9 along x-axis denote the study groups.

**Figure 12:**
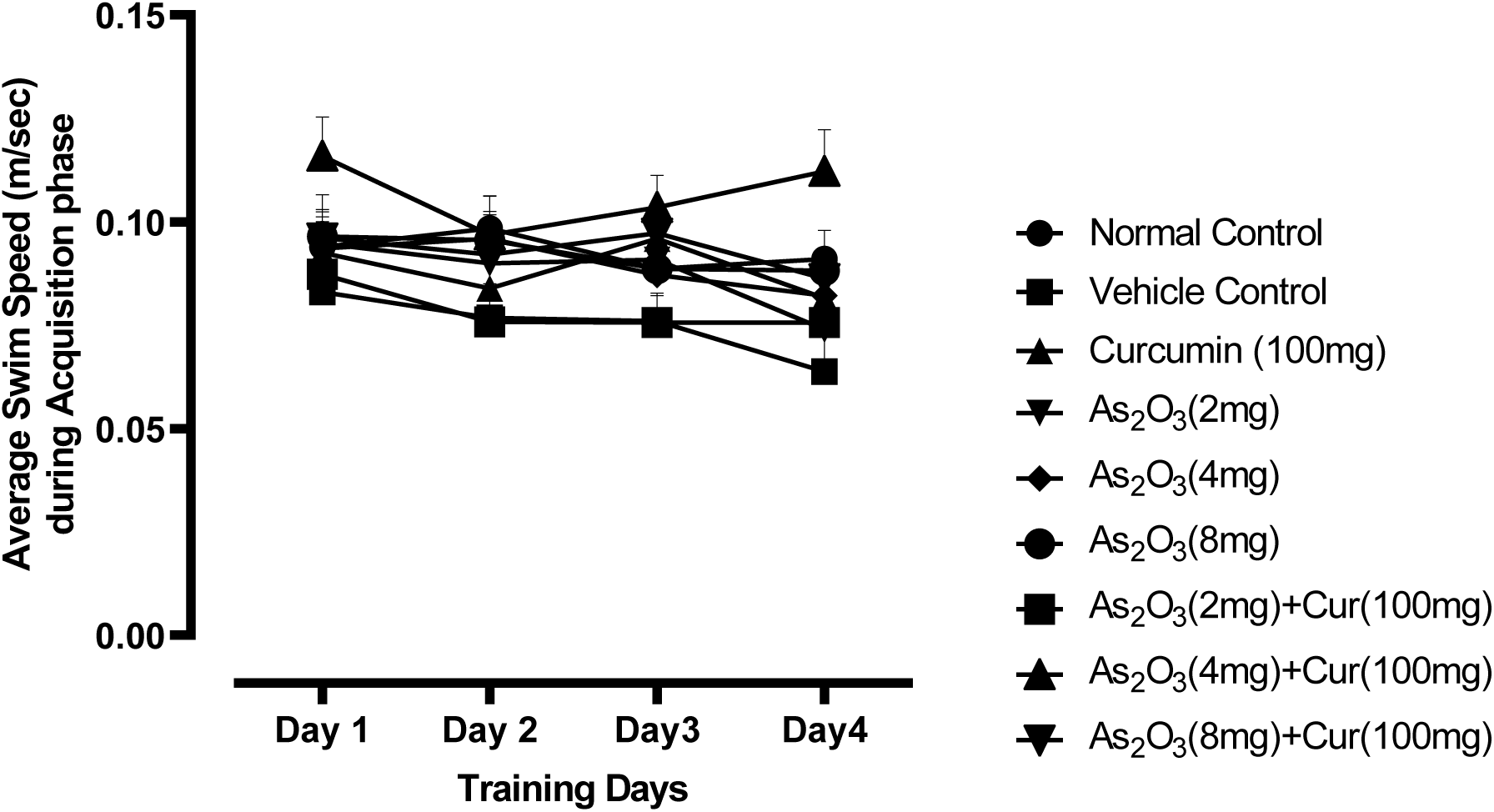
The average swim speed during acquisition phase of MWM test (n=12).

### Probe trial

The reference memory was assessed 24 hours following the last day of acquisition phase by performing a probe trial. The preference for platform area was determined by recording the number of platform area crossovers and the time spent in the target quadrant by the animal.

The association between the number of platform crossovers and the animal groups was not found to be significant (p<0.0486) as shown in Figure 13A & S13. It was observed that the median values for the number of platform crossovers were lesser in As_2_O_3_ alone treated groups (2, 4 & 8mg/kg) as compared to the control & As_2_O_3_ + CUR co-treated groups (2, 4 & 8 mg/kg + CUR 100 mg/kg).

**Figure 13:**
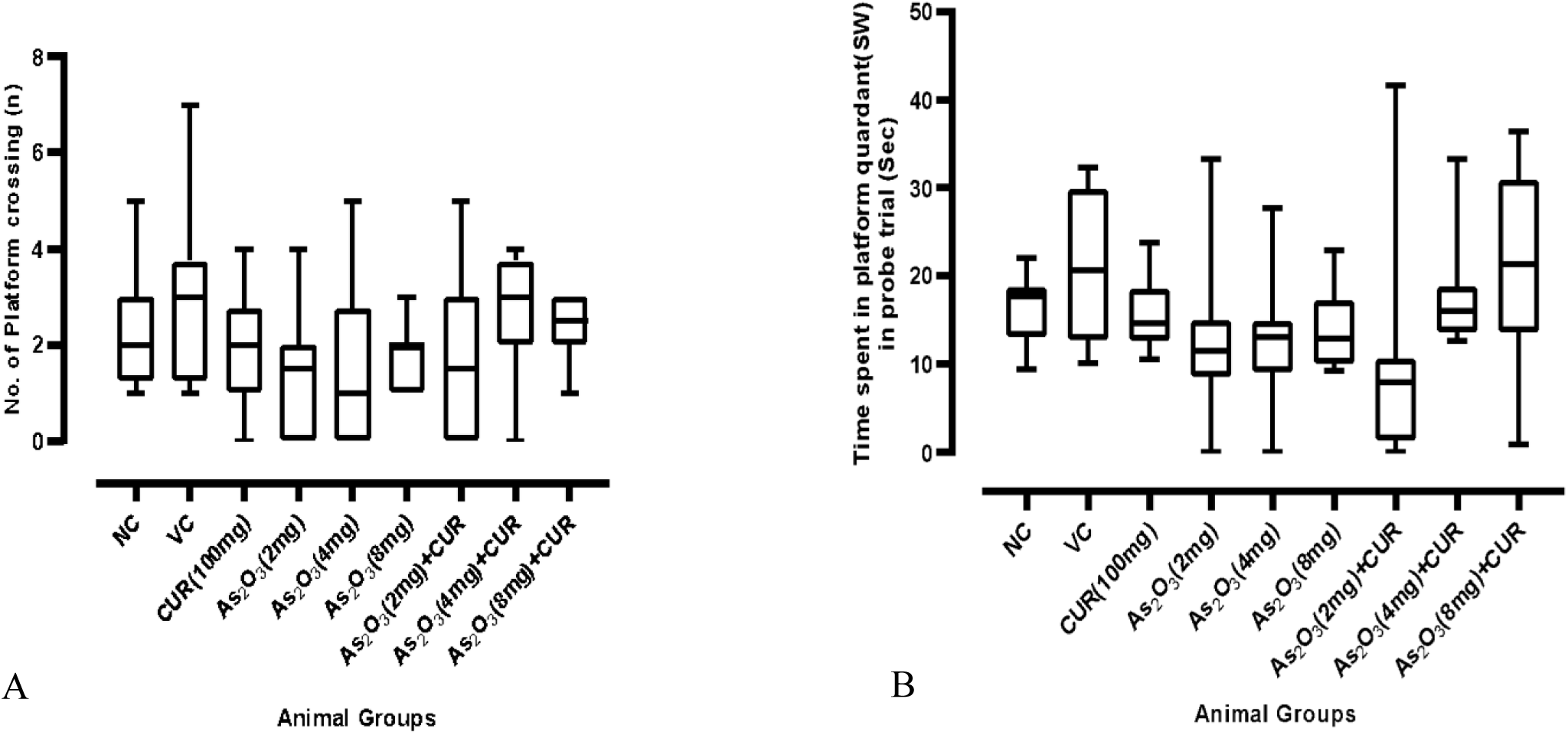
(A) Number of platforms crossing (n) and (B) time spent in platform quadrant (sec) in MWM test during probe trial day among control and experimental groups (n=12 animals/group).

The time spent in the target quadrant was found to be significant across the groups (p<0.0002) (Figure 13B & S13). The mean percentage time spent in the target quadrant was longer for the animals which received CUR with As_2_O_3_ (4 & 8 mg/kg) as compared to the As_2_O_3_ alone treated groups (4 & 8 mg/kg) and found to reach the significance levels (p<0.0410, p<.0433).

MWM observations do suggest that CUR administration along with As_2_O_3_ apparently enhances the learning ability and retention memory in comparison to As_2_O_3_ alone treated groups. However, a higher dose of CUR over a more prolonged period might bring out changes of statistical significance.

### 3.3 Effects of CUR supplementation on basal ganglia-mediated biochemical functions on exposed mice

#### 3.3.1 Reduced Glutathione (GSH)

A dose dependent significant decrease (p<0.0009) was observed in GSH levels of animals receiving As_2_O_3_ 2, 4 & 8 mg/kg alone with the median values 1.67(0.91-4.63), 1.81(0.18-3.59), 0.55(0.32-0.96) respectively as compared to the control with the median value 4.62(2.15-7.2) (Figure 14A & S14). As_2_O_3_ 2,4 & 8mg/kg alone treated mice showed a significant down regulation of GSH relative to control mice (p<0.0247, p<0.0104, p<0.0038) respectively, whereas co-administration of CUR along with As_2_O_3_ (2, 4 & 8 mg/kg) with the median value 2.06(1.1-4.58), 2.95(2.52-3.57), 4.71(1-5.55) respectively, resulted in up regulation of GSH levels as compared to As_2_O_3_ 2mg, 4mg & 8 mg/kg alone treated with the median value 1.67(0.91-4.63), 1.81(0.18-3.59), 0.55(0.32-0.96) respectively, out of which As_2_O_3_ 8 mg/kg alone treated with the median value 0.55(0.32-0.96) showed significant decrease (p<0.0038) level of GSH as compared to co-administration of CUR along with As_2_O_3_ 8 mg/kg with the median value 4.71(1-5.55), thereby suggestive of Curcumin-induced up regulation of GSH levels in basal ganglia of mice.

**Figure 14:**
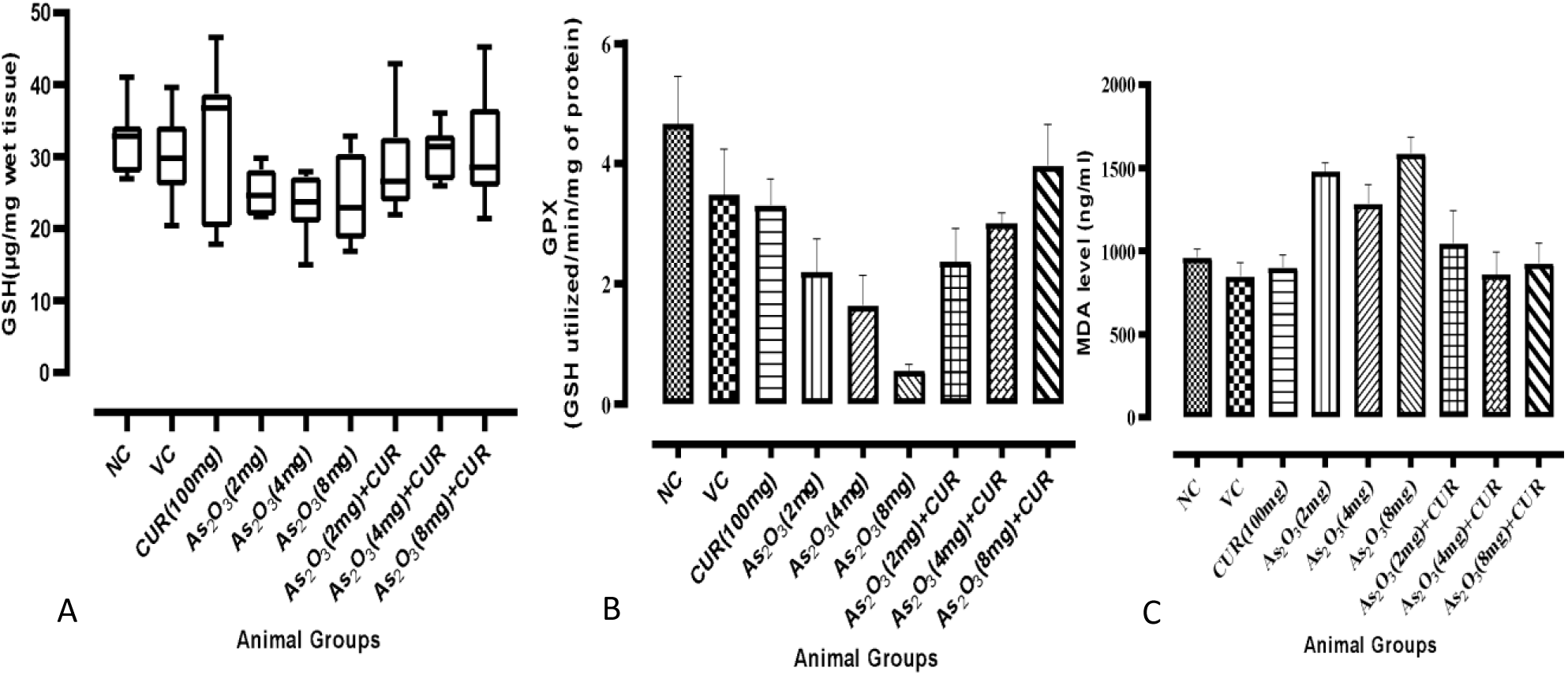
GSH levels (A), GPX (B) and Lipid peroxidation level MDA (C) in various control and experimental in striatal tissue (n=7 animals/group).

#### 3.3.2 Glutathione peroxidase (GPx)

Similar pattern was observed in the level of glutathione peroxidase in the basal ganglia region but none of them showed significant changes (p<0.0346). Apparently, decreased GPx levels were found in basal ganglia of mice treated with As_2_O_3_ 2, 4 & 8 mg/kg alone with the mean values 25.04 ± 3.163442, 23.07571 ± 4.437638, 23.84286 ± 5.856622 respectively as compared to the controls with the mean value 32.50143 ± 4.732365 (Figure 14B & S14). Whereas co-administration of CUR with As_2_O_3_ (2, 4 & 8 mg/kg) showed mean values of 29.20286 ± 7.101879, 30.91286 ± 3.537493, 30.66857 ± 7.859117 respectively and led to increased GPx levels as compared with As_2_O_3_ 2mg, 4mg & 8 mg/kg alone treated group with the mean values of 25.04 ± 3.163442, 23.07571 ± 4.43763 & 23.84286 ± 5.856622 respectively.

#### 3.3.3 Malondialdehyde (MDA)

The mean MDA levels ranging from minimum 859.61 ± 327.33 to maximum 1587.34 ± 238.15, were found to be comparable across the groups (P<.001) and the values are shown in figure 14C & S14. MDA levels in animals receiving As_2_O_3_ 2, 4 & 8 mg/kg alone were 1478.84±129.62, 1285.41±284.44, 1587.34±238.15 (mean±SD) respectively as compared to the control group with the mean value of 961.85 ±118.68. The difference between control & As_2_O_3_ 8 mg/kg alone treated groups were found to be statistically significant (P<.009). Co-administration of CUR along with 2mg, 4mg, & 8mg/kg As_2_O_3_ with the mean value 1044.55 ± 481.78, 859.61 ± 327.33, 926.76 ±

297.05 respectively found to decreased MDA levels as compared to As_2_O_3_ 2mg, 4mg & 8 mg/kg alone with the mean value 1478.84 ± 129.62, 1285.41 ± 284.44, 1587.34 ± 238.15 respectively, out of which there is also observed a significant increased (P<.005) MDA level in 8mg/kg As_2_O_3_ with the mean value 1587.34 ± 238.15 alone treated group as compared to co-administration of CUR along with 8mg/kg As_2_O_3_ with the mean value 926.76 ± 297.05. Together, the observations of biochemical study were suggestive of Curcumin-induced up regulation of antioxidant enzymes (GSH & GPx) along with down regulation of oxidative stress markers (MDA) in the basal ganglia region of mice.

### 3.4 Morphometric observations in Caudate Putamen

The CV stained 7µm coronal sections (paraplast) (at Bregma level +0.98mm to +0.02mm) of mouse brain (control and experimental animals) were observed under light microscope for morphological features of various regions of caudate putamen (CPu) nucleus (DM, DL, VM & VL) based on their anatomical orientation. We have observed the area of neurons as well as neuronal density in above mentioned regions of CPu as shown in figure 15.

**Figure 15:**
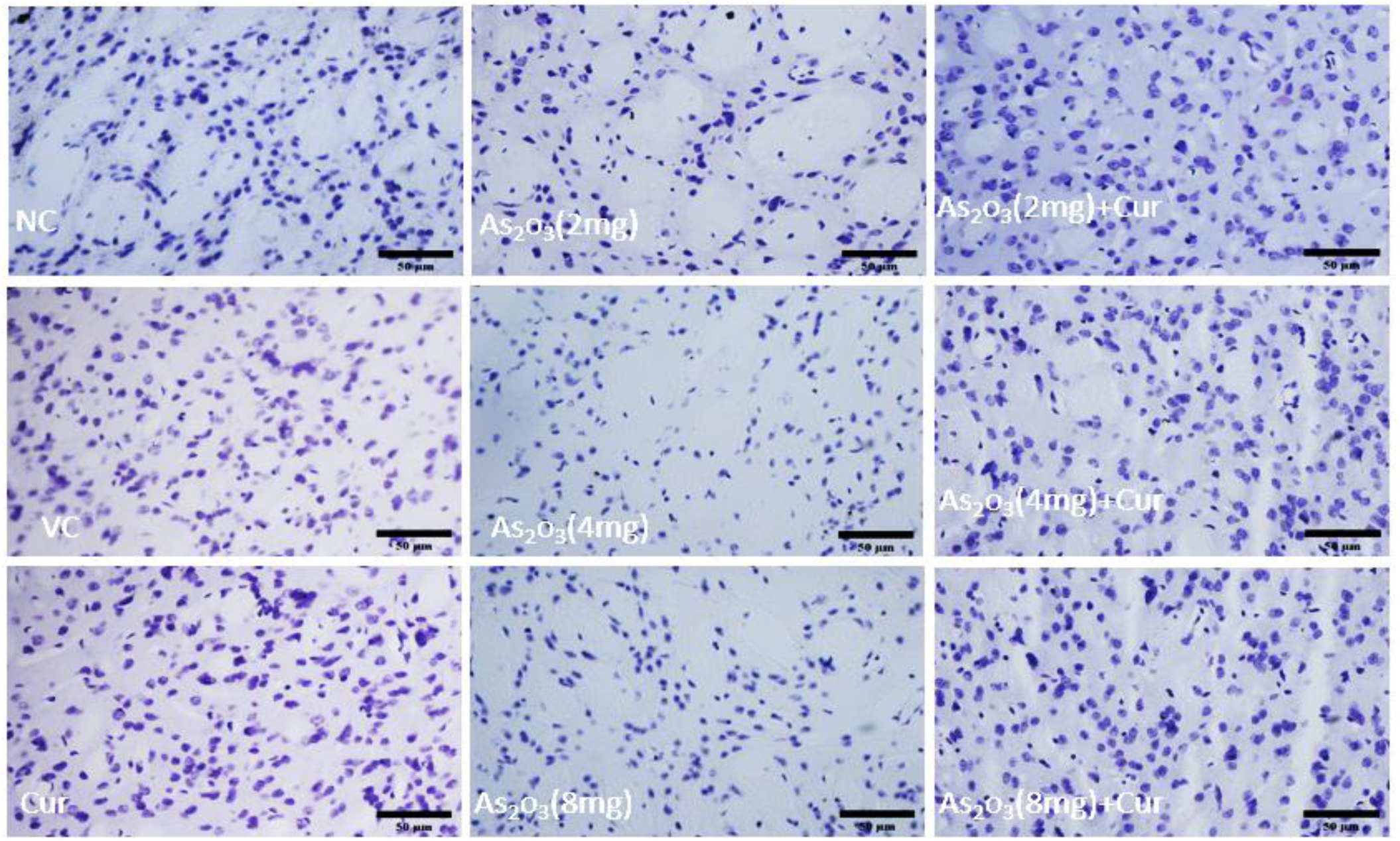
High magnification (40X) photomicrographs showing CV staining in coronal sections of Caudate putamen of control and experimental groups (n=6)

#### 3.4.1 Neuronal area in Dorsal & Ventral Striatum

In our observation the mean neuronal area in dosrsal striatum is ranged from minimum 51.57±6.34 µm^2^ to maximum 63.95±10.86 µm^2^ also comparable across the groups (P<.0513) and the values are showing in figure 16A & S15. It is observed that neuronal area in found to be decreased in 4mg & 8mg As_2_O_3_ with the mean value 51.57 ± 6.34 µm^2^, 51.71 ± 6.43 µm^2^ respectively, as compared to control group with the mean value of 52.96 ± 6.87 µm^2^. Whereas co-administration of CUR along with As_2_O_3_ (4 & 8 mg/kg) with the mean value 59.78 ± 8.64 µm^2^, 61 ± 10.9 µm^2^ respectively, led to increase in the dorsal striatum neuronal area as compare with As_2_O_3_ 4mg & 8 mg/kg alone treated group with the mean value 51.57 ± 6.34 µm^2^, 51.71± 6.43 µm^2^ respectively. However, it was not found to be statistically significant (p>0.05).

**Figure 16:**
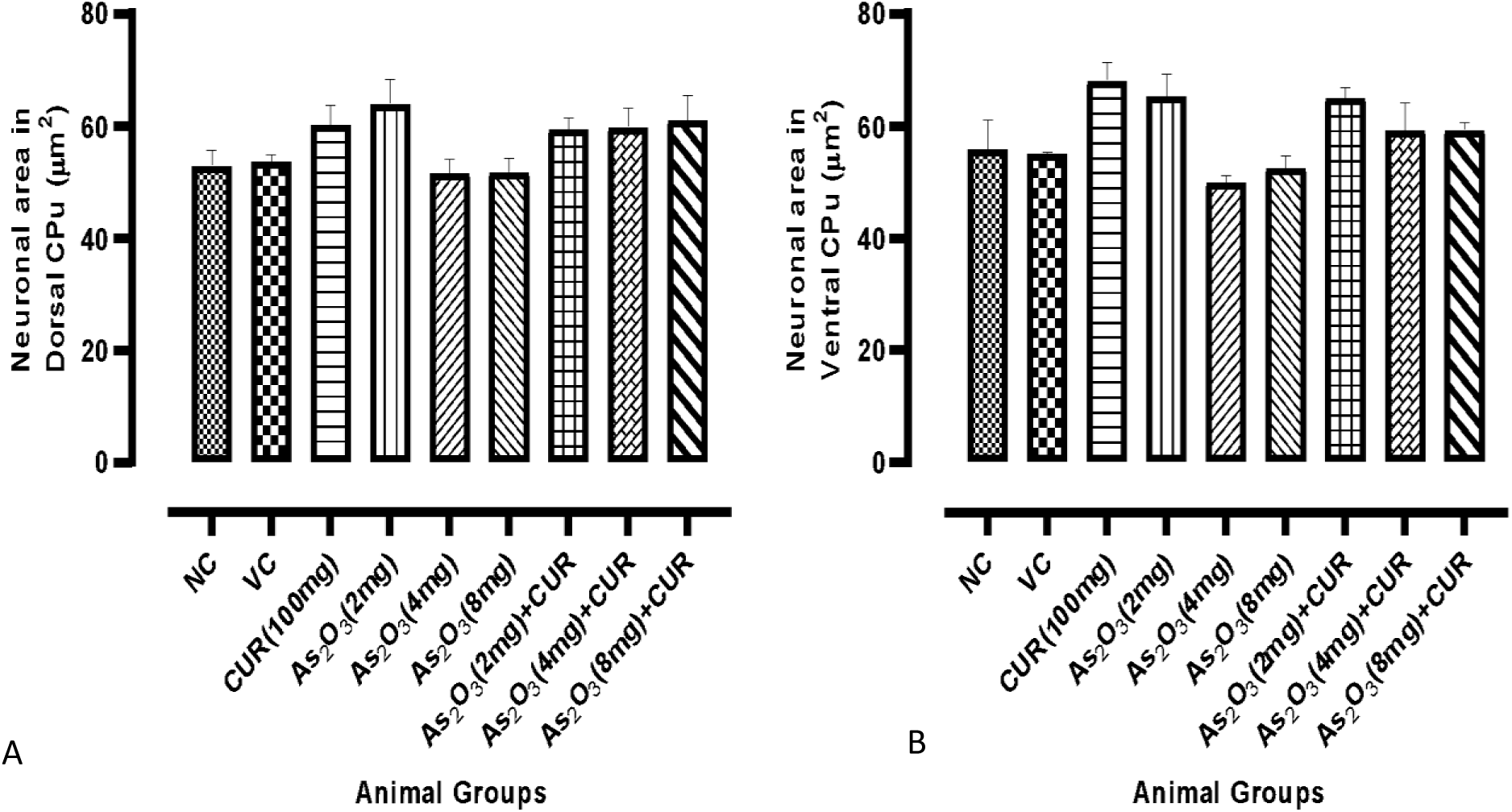
The neuronal area in (A) dorsal and (B) ventral caudate putamen (CPu) region of control and experimental groups (n=6 animals/group).

The mean ventral striatal neuronal area ranged from minimum 49.95 ± 3.07µm^2^ to maximum 68.12 ± 7.88µm^2^ also comparable across the groups (P<.0018) and the values are shown in 16B & S15. It was observed that neuronal area was decreased in 4 & 8mg of As_2_O_3_ with the mean values 49.95 ± 3.07 µm^2^, 52.52 ± 5.32 µm^2^ respectively, as compared to control group with the mean value of 55.79 ± 13.09 µm^2^. Whereas Co-administration of CUR along with As_2_O_3_ (4 & 8 mg/kg) with the mean values 59.25 ± 12.1 µm^2^, 59.29 ± 3.36 µm^2^ respectively, led to increase in the neuronal area of dorsal striatum as compare with As_2_O_3_ 4mg & 8 mg/kg alone treated group with the mean value 49.95 ± 3.07 µm^2^, 52.52 ± 5.32 µm^2^ respectively. However, it was not found to be statistically significant (p>0.05).

#### 3.4.2 Density of neurons in dorsal & ventral striatum

In our observation the median neuronal density in dorsal striatum is ranged from minimum 50.62(28.5-77.75) to maximum 63.12(13.75 - 124) also comparable across the groups (P<.0153) and the values are showing in figure 17A & S16. It is observed that mean neuronal density found to be decreased in 2mg, 4mg & 8mg As_2_O_3_ with the median value 54.62(21.5-59.5), 54.87(43.75-57.5), 54.87(43.75-57.5) in numbers respectively, as compared to control group with the median value of 63.12(13.75 - 124). Whereas co-administration of CUR along with As_2_O_3_ (2 mg/kg) with the median value 58.75(52-61) observed the increase in the dorsal striatum neuronal cell density as compare with As_2_O_3_ 2mg/kg alone treated group with the mean value 54.62(21.5-59.5). However, it does not reach significant level (p>0.05).

**Figure 17:**
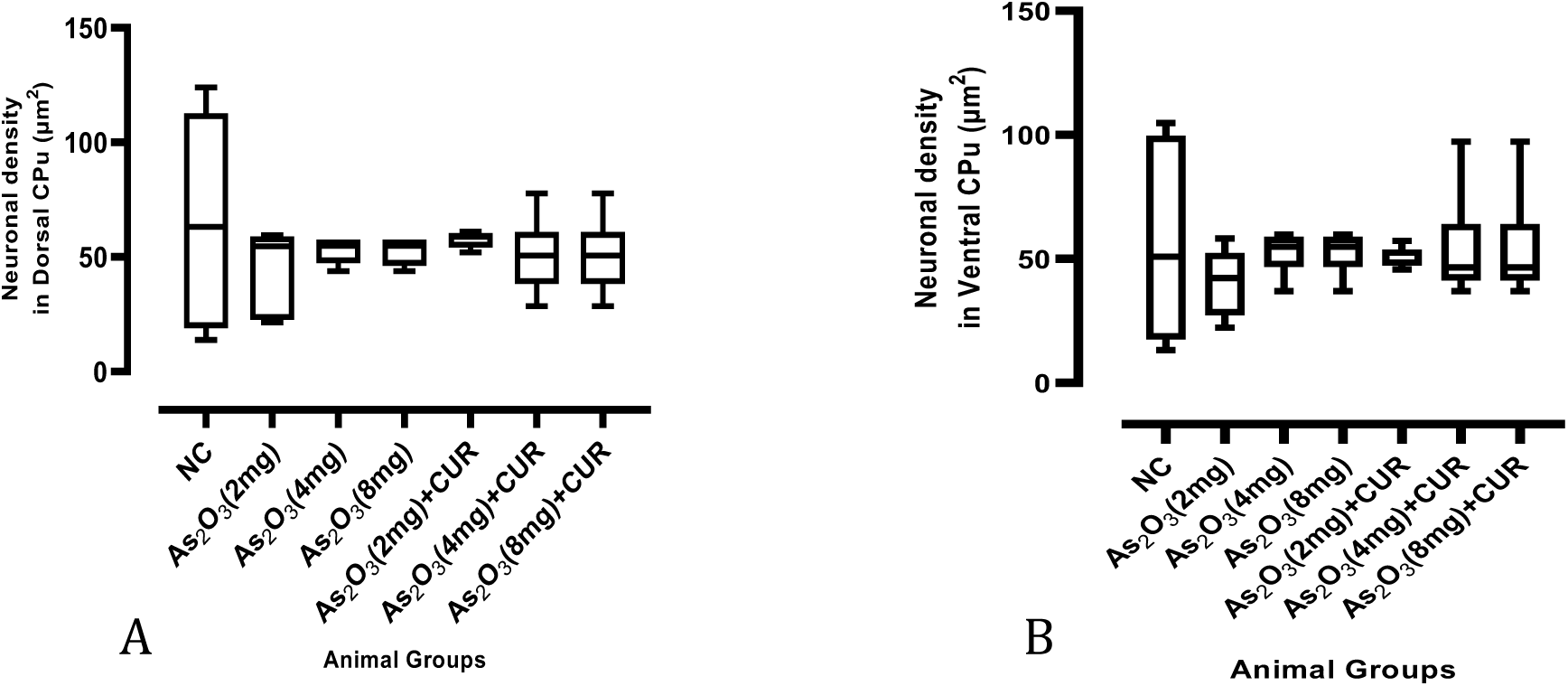
The neuronal density in (A) dorsal and (B) ventral caudate putamen (CPu) region of control and experimental groups (n=6 animals/group).

Median ventral striatal neuronal cell density ranged from minimum 42.25(22.25 - 58.25) to maximum 54.87(37 - 59.75) in number, also comparable across the groups (P<.0051) and the values are showing in figure 17B & S16. It is observed that median neuronal cell density was found to be decreased in 2mg/kg As_2_O_3_ treated with the median value 42.25(22.25 - 58.25) as compared to control group with the mean value of 50.87 (1.25-104.75). Whereas Co-administration of CUR along with As_2_O_3_ 2mg/kg with the median value 48.5(45.75 - 57.25) led to increase in the dorsal striatum neuronal cell density as compare with As_2_O_3_ 2mg/kg alone treated group with the mean value 42.25(22.25 - 58.25) in number. However, it was not found to be statistically significant (p>0.05).

## 4. Discussion

The morbidity due to induction of long-term exposure of *As* is a major health concern because of its bioaccumulation and non-biodegradation in organs like liver, kidneys, lungs, brain etc. [47,48]. With context to nervous system, chronic *iAs* exposure has been reported to induce memory decline, decrease in speed of information processing, anxiety, cognitive dysfunction, and depression-like behaviors [49,50]. A dose dependent effect of *iAs* in animal models of neurotoxicity has been previously reported [51,52]. However, there is no single active agent that has been identified as an antidote against *iAs*-induced toxicity [53]. Due to the association between neurotoxicity and oxidative stress caused by exposure to *iAs*, phytochemicals can be identified as effective preventive or therapeutic agents against *iAs*-induced toxicity, which is also being recommended by WHO [54].

Hence, the current study focused on the effects of exogenously administered Curcumin (antioxidant) on behavioral, biochemical, morphological and morphometric parameters with context to basal ganglia of mice subjected chronically to As_2_O_3_.

In the present study, there was no significant alteration observed in the body weight of animals across the groups at the end of the experimental period as observed by studies of former investigators [55,56]. Our observations with no significant alterations in body weight of animals across the groups could be attributed to shorter duration of exposure to As_2_O_3_, which might not have been sufficient to influence the body weight.

The general physical features (body weight, grooming, rearing, state of alertness, etc.) were maintained during the experimental period across all the groups. However, Transitory hair loss (patchy) was observed in animals receiving higher dose of As_2_O_3_ (8 mg/kg) as reported by earlier studies [15,57]. No significant dose effect was observed on the general features and percentage change in the body weight of As_2_O_3_ alone exposed animals when compared to controls and CUR co-treated groups.

Earlier studies have reported anxiety-like behavior in mice and psychiatric frailty, i.e., anxiety and depressive disorder following *iAs* exposure [58]. Hence, we conducted Open field test (OFT) to assess the exploratory and locomotor activity along with anxiety status in rodents subjected to exogenously administered substances [59].

The present study showed no significant alteration observed in exploratory and locomotor activity of experimental animals (exposed to As_2_O_3_ alone or As_2_O_3_+ CUR) during OFT testing. The animals receiving a higher dose (8 mg/kg) of As_2_O_3_ showed less time spent in the central zone and increased time spent in the peripheral zone, suggesting thereby apparent increase in anxiety levels. Exposure of animals to 8 mg/kg As_2_O_3_ also resulted in anxiety-like behavior and manifested in the form of a significant decrease in the number of center entries and an increase in the number of corner entries.

Various neurochemical findings have been linked to dopaminergic hypo function induced by *iAs* exposure [60]. The dopaminergic system is reported to be extremely sensitive to oxidative stress (OS) and the addition to the free radicals generated by *iAs* toxicity increases the level of OS [61]. Kumar and Reddy observed significant impairment in OF tasks including number of line crossings, rearing, grooming episodes in rats exposed to NaAsO_2_ (10 mg/kg bw oral gavage) suggesting thereby NaAsO_2_ induced adverse impact on locomotor and exploratory activity along with increases in anxiety levels [49]. Adedara and co-worker exposed rats to 60µg NaAsO_2_/L alone or in combination with inorganic and organic forms of selenium (Se - 0.25 mg/kg; diphenyl diselenide (DPDS) (2.5 mg/kg) for 45 successive days [50]. NaAsO_2_ alone induced rats in OFT showed marked impairment in the locomotor activity and anxio-genic like behavior in the form of decreased average speed, total distance travelled in OFT. The supplementation of Se (inorganic or organic form) with NaAsO_2_ augmented the exploratory activity in rats and modified the status of endogenous antioxidant levels such as reduced glutathione (GSH) levels in discrete organs (cerebellum, cerebrum, and liver) and decreased levels of AChE activity (cerebellum and cerebrum). The use of natural antioxidants such as CUR has been studied by various investigators working on *iAs* toxicity to evaluate its role in restoration of arsenic-induced neurobehavioral alterations. Anxiolytic effect of CUR has been reported based on OFT parameters studied in several studies [62,63].

Abu-Taweel et al. reported significant protective effect of CUR (150 & 300 mg/kg bw) on cadmium (Cd) induced adverse effects on body weight, anxiety status, learning capability (cognitive effect) and muscular activity in Swiss-Webster strain male mice [64]. Wang et al. demonstrated that pre-treatment with CUR (50 mg/kg) for 7 days reversed the depression-like behavior in LPS-induced mice model of depression [65]. Haider et al. reported the prevention of both anxiety (EPM) and hyperactivity (OFT) produced in response to acute immobilization stress in rats when pre-treatment with CUR was carried out [63].

Previous investigations have pointed towards dose and duration of *iAs* exposure as important determinants in neurobehavioral alterations [15]. *iAs*-induced apoptotic neuronal damage in certain brain regions such as cortex & hippocampus could be another possibility of development of anxiety-like behavior in rodents [19]. Based on region-specific altered expression of various agents involved in antioxidant and dopaminergic systems, Rodriguez and coworkers observed that low levels of *iAs* exposure could cause subtle changes in nervous system, but manifestation of behavioral changes becomes evident only after exposure to high dose *iAs* [15].

Various studies have been reported that exposure of *As* in humans causes central and peripheral neuropathy [66]. Rodriguez and co-workers observed that dose, duration, route of exposure are the most important factors underlying *iAs*-induced altered locomotor activity [51]. In the present study, we have observed that the median latency to fall in rota-rod experiments was comparable across the groups. We have also found that As_2_O_3_ alone exposed groups showed significant reduction in latency to fall, when compared to control groups. Whereas co-administration of As_2_O_3_+CUR treatment was shown to considerably increase the latency to fall when compared with As_2_O_3_ alone exposed groups but did not reach the significance level. *As*-exposure is associated with changes in the Rota-rod performance of mice as they fell early from the rotating rod, thereby suggesting impairment of motor co-ordination. Curcumin (CUR) treatment in *As* exposed mice protected the ability of the *As-*induced changes in motor co-ordinations.

Next, we have determined object recognition memory in mice exposed to As_2_O_3_ by NOR task. In our study, we noticed that a reduction in the memory index, resulting from exposure to *iAs*, indicated an impairment in learning and memory formation in mice. However, when mice exposed to *iAs* were treated with curcumin (CUR), the memory index closely resembled that of the control groups. These results imply that co-administration of CUR effectively prevented the learning and memory impairment induced by *iAs* exposure. In line with our findings, previous studies have shown the memory impairment responses in NOR experiments following cadmium (Cd) exposure [67–70]. Previous reports have shown that CUR administration is able to attenuate the oxidative damage and prevent memory loss in rats exposed to other neurotoxicity chemicals, such as aluminum chloride [71], arsenic [22], and cigarette smoke [72].

The Morris Water Maze (MWM) test is a well-established animal behavioral test for the assessment of learning and memory. In the present study, a significant fall in EL on day 4, when compared to day 1 during acquisition phase, was observed in control and As_2_O_3_ + CUR co-treated groups, reflecting normal learning ability. The difference in EL on day 4 as compared to EL on day 1 was not significant in As_2_O_3_ alone induced groups as compared to control and As_2_O_3_ + CUR co-treated groups. In the probe trial, the number of platform crossovers and time spent in the target quadrant was apparently lesser in As_2_O_3_ alone induced groups than the control and As_2_O_3_ + CUR co-treated groups. These observations align with and support the findings documented in previous studies [73–75].

Jiang et al. (2014) exposed weaning male SD rats to sodium arsenite (5.5 and 8.2 mg/kg bw) for 3 months and observed impaired performance in the MWM test in exposed groups, with the degree of impairment varying with the arsenic dose. Control mice in the hidden platform test displayed a progressive decline in escape latencies and path lengths over 8 training days, relevant to findings reported by Krüger et al. (2006), who noted inhibition of long-term potentiation after arsenite exposure. Additionally, impairment of conscious learning has been linked to impaired LTP. Based on these findings, Jing and colleagues suggested that inhibition of LTP induction following arsenic exposure might be a contributing factor to impaired conscious learning.

Previous research has shown that CUR can enhance learning and memory by harnessing its antioxidant properties. Oz et al. (2015) found that CUR treatment reversed cognitive impairment caused by cisplatin, as evidenced by increased time spent in the target quadrant and more crossings over the original platform area in the chronic CUR supplementation group compared to the cisplatin-treated group [4]. They also noted that CUR restored MDA, SOD, and AChE activities, suggesting that CUR enhances cholinergic function and exerts antioxidant effects. These observations do substantiate the findings of the present study, whereby the apparent alterations observed in spatial learning and memory (MWM) induced by As_2_O_3_ alone exposure were improved substantially in animal groups receiving CUR supplementation along with As_2_O_3_.

Several studies have documented the role of arsenic as a pro-oxidant agent inducing OS in various body tissues [76–78]. Hence, we determined the levels of antioxidant enzymes and lipid peroxidation. In the present study, increased levels of MDA as well as decrease in antioxidant levels (GSH & GPx) was observed in basal ganglia of As_2_O_3_ alone treated groups compared to the controls, and the level of increase was more pronounced with higher dosages of As_2_O_3_ (4 & 8 mg/kg bw). These findings agree with the observations of previous studies pertaining to heavy metal toxicity [79] and use of As_2_O_3_ as a potent chemotherapeutic agent [80]. Increase in MDA level is also associated to decreased activity of antioxidant enzymes like catalase (CAT) and glutathione peroxidase (GPx) [49,77], which normally reduces H_2_O_2_ into water and nascent oxygen (O). *As* is presumed to have a direct effect on the protein structure of GPx causing its complete inhibition or loss of activity. Increased H_2_O_2_ may result in cell membrane damage as H_2_O_2_ derived hydroxyl radical (OH) may cause hydrogen abstraction from the cell membrane [77].

In the present work, the reduction in GSH level observed in As_2_O_3_ alone treated groups was more pronounced with higher dosages of As_2_O_3_ (4 & 8 mg/kg). These findings are in concordance with the earlier reports [75,77,81,82]. Depletion of GSH may be due to its utilization either by the toxic radicals or by the intermediates formed during methylation of arsenic in presence of SAM (S-adenosylmethionine) [82]. Another reason for low levels of GSH may be the direct binding of arsenic with thiol groups present in GSH to form As(GS)_3_ [83–85].

Various reports regarding ineffectiveness of chelating agents against arsenic induced toxicity led to the trial of natural antioxidants like CUR to counter the arsenic-induced OS. In the present study, the anti-oxidant activity of CUR was advocated by decreased levels of MDA and increase in GSH & GPx level in groups that received CUR supplementation along with As_2_O_3_. CUR is labeled as a potent neuroprotectant due to its effective antioxidant property. Its free radical scavenging activity is considered to be due to its poly phenolic structure as compared to single phenol hydroxyl group present in other flavonoids [86,87] and also the number of –CH_2_ sites it possesses. In addition, a number of studies carried out in various animal models and humans have demonstrated the extreme safe nature of CUR even at very high doses [88]. Neuroprotective efficacy of CUR has been studied with context to its varied range of doses (40 to 200 mg/kg body weight), different time durations and multiple routes of exposure in animal models of diseased conditions and neurotoxicity [89,90]. The ability of CUR to cross BBB [91,92] provides further supportive evidence to its neuroprotective potential [93,94]. The ability of CUR to inhibit the generation of superoxide radicals by the increased production of GSH (a major non-protein thiol in the living organism) does play a critical role in maintaining body’s antioxidant defense system [14,87,95]. It may be reinforced that CUR counteracts the arsenic induced tissue damage by quenching the generation of ROS.

In the present study, the observation of the shrunken neurons and spaces here and there in the caudate putamen region of As_2_O_3_ alone exposed groups are in accordance with the earlier reports where arsenic-induced morphological changes in brain sections of rodent models have been put forth. These observations do suggest that neurons could be one of the major target cells of arsenic neurotoxicity. On the other hand, the neuronal morphology in As_2_O_3_ + CUR co-treated groups was comparable to controls. Yang et al. (2005) in their in-vitro study on primary rat hippocampal neurons reported arsenic-induced apoptotic features such as decrease in viable cell growth, cytoplasmic vacuoles, frothing and nuclear condensation with intact cell membrane [91]. Chattopadhyay and co-worker carried out their experimental study on natal mice and observed *iAs-* induced abnormalities in neuronal cell membrane, loss of ground matrix, inhibited neural networking and apoptosis [96].

Earlier studies reported vacuolar degeneration in cytoplasm along with karyorrhexis and karyolysis [97] in brain sections of mice exposed to varying doses of As_2_O_3_ (4 mg/L; 1ppm; 2ppm) for 15 days. These investigators further noted mild histo pathological changes in animals receiving As_2_O_3_ + taurine suggesting the positive influence of taurine on As2O3-induced adverse effects. Decrease in neuronal density and their deranged arrangement in association with karyopyknosis & karyolysis was reported in the CA1 area of hippocampus of rats exposed to 4 ppm As_2_O_3._ Somewhat similar neuronal features such as edematous changes and degeneration along with focal gliosis have been described by various investigators in brain sections of animal models following arsenic exposure [81,98]. These morphological alterations changes have been associated with oxidative stress and initiation of apoptotic cascade induced by toxic chemicals.

Fallah et al. (2018) investigated the effect of CUR and NAC (N-acetyl-cysteine) on brain histology in rats exposed to 20 mg/kg arsenic through oral gavage (every other day for 30 days) and observed cellular damage and severe lesions in white matter of hippocampus and CA1 neurons in arsenic alone exposed rats, whereas normal gray and white matter along with well-arranged neurons with normal density and morphology were found in rats co-treated either with CUR or NAC [78]. Also, the antioxidant treated groups showed neuronal count near to that observed in the control group. These findings do suggest the protective effect of CUR and NAC against arsenic –induced toxicity. Adedara et al. (2020) reported marked neuronal degeneration in rat brain and morphological changes in liver following *iAs* intoxication (60 µg NaAsO_2_/L) [50]. In their detailed report, these investigators correlate various morphological alterations with induction of caspase-3 along with oxidative and inflammatory stress induced by arsenic exposure.

In conclusion, based on the present findings, it becomes evident that As_2_O_3_ exposure adversely impacts the behavioral, morphological and biochemical parameters with context to basal ganglia in a dose dependent manner and exogenous supplementation of CUR (100 mg/kg) proves to be beneficial in amelioration of these As_2_O_3_-induced adverse effects to some extent.

## 5. Future perspectives

Curcumin has demonstrated an attenuating effect on arsenic induced neurotoxicity in mice. Looking ahead, further comprehensive research is required to understand the long-lasting effects of *iAs* exposure on the expression levels of various key neurotransmitters which play a critical role in the functioning of basal ganglia. The expression of the human genome is a complex process and is regulated by multiple factors. The transcriptomics and proteomics analysis of differentially expressed genes is required to understand the exact mechanisms of Arsenic toxicity and the genes involved in Neurodegeneration induced by *iAs* exposure. The detailed analysis of transcriptomics will provide the driver genes involved in *As-*induced toxicity which will facilitate the strategic selection of novel therapeutic agents to combine with *As* treatment in order to reduce its toxic impact while maintaining the anti-cancerous activity.

