## Supplementary tables for "Curcumin alleviates Arsenic trioxide-induced behavioural impairment, oxidative damage & morphological alterations in striatal region of mice brain": Supplementary Tables.pdf

**S1:** Descriptive statistics of percentage body weight (grams) change from Day 1 to 45 among control and experimental animals.

| Groups | Body Weight (g) mean ± SD | | |
|---|---|---|---|
| | **Day1** | **Day45** | **Percentage Change** |
| Normal Control (NC) | 24.08±.46 | 39.18±2.82 | 62.71±11.54 |
| Vehicle Control (VC) | 23.57±.46 | 38.8±4.17 | 64.58±17.37 |
| Curcumin(100mg) | 23.45±.45 | 37.25±3.25 | 58.62±13.36 |
| $As_2O_3$(2mg) | 22.71±.89 | 41.99±2.85 | 85.10±14.85 |
| $As_2O_3$(4mg) | 23.90±.56 | 38.4±4.79 | 60.60±19.94 |
| $As_2O_3$(8mg) | 23.85±.66 | 37.79±3.77 | 58.73±18.12 |
| $As_2O_3$(2mg)+Cur(100mg) | 22.79±1.11 | 38.90±4.41 | 70.88±19.93 |
| $As_2O_3$(4mg)+Cur(100mg) | 22.79±.76 | 37.65±4.15 | 65.14±16.66 |
| $As_2O_3$(8mg)+Cur(100mg) | 22.73±.77 | 40.11±2.91 | 76.73±15.43 |
| ***p*-Value** | ***p*<.024** | ***p*<.542** | ***p*<.713** |

Values are expressed as Mean ± SD (n=12), $p < 0.05$ compared with control.

**S2:** Tabulated data of velocity (m/sec) of control and experimental animals in OFT.

| **Groups** | **Velocity** | ***p* value** |
|---|---|---|
| Normal Control (NC) | 0.025 (0.01 - 0.05) | a) $p < 1.000$<br>b) $p < .450$<br>c) $p < .071$<br>d) $p < 1.000$<br>e) $p < 1.000$<br>f) $p < .781$ |
| Vehicle Control (VC) | 0.03(0.01 - 0.05) | |
| Curcumin(100mg) | 0.02(0.01 - 0.04) | |
| $As_2O_3$(2mg) | 0.02(0 - 0.04) | |
| $As_2O_3$(4mg) | 0.02(0 - 0.03) | |
| $As_2O_3$(8mg) | 0.01(0.01 - 0.03) | |
| $As_2O_3$(2mg)+Cur(100mg) | 0.02(0.01 - 0.04) | |
| $As_2O_3$(4mg)+Cur(100mg) | 0.02(0.02 - 0.03) | |
| $As_2O_3$(8mg)+Cur(100mg) | 0.025(0.01 - 0.03) | |
| **Overall ANOVA *p*< .0231** | | |

Data presented as median (min-max) values. The results were analysed using Bartlett's test for equal variance of One-way ANOVA followed by Bonferroni post-hoc test for multiple comparison between the groups. Number of pairwise comparisons denoted in the table are as follows; (a) NC vs $As_2O_3$(2mg) alone; (b) NC vs $As_2O_3$(4 mg) alone; (c) NC vs $As_2O_3$(8 mg) alone; (d) $As_2O_3$(2mg) alone vs $As_2O_3$(2mg)+CUR; (e) $As_2O_3$(4mg) alone vs $As_2O_3$(4mg)+CUR; (f) $As_2O_3$(8mg) alone vs $As_2O_3$(8mg)+CUR.

**S3:** Descriptive statistics of line crossing (n) of control and experimental animals in OFT.

| **Groups** | **Line crossing** | ***p* value** |
|---|---|---|
| Normal Control | 69.25 (26 - 113) | a) $p< 1.000$<br>b) $p< .012$<br>c) $p< .039$<br>d) $p< 1.000$<br>e) $p< 1.000$<br>f) $p< 1.000$ |
| Vehicle Control | 60(21.5 - 117) | |
| Curcumin(100mg) | 61.75(31 - 102) | |
| $As_2O_3$(2mg) | 62.5(22 - 95.5) | |
| $As_2O_3$(4mg) | 34.75(2 - 66.5) | |
| $As_2O_3$(8mg) | 30 (18 - 99.5) | |
| $As_2O_3$(2mg)+Cur(100mg) | 47.75(20 - 102) | |
| $As_2O_3$(4mg)+Cur(100mg) | 43.75(30 - 66.5) | |
| $As_2O_3$(8mg)+Cur(100mg) | 35(19 - 67.5) | |
| **Overall ANOVA *p*< .0004** | | |

Data expressed as median (min-max) (n=12/group). Result analysed using Bartlett's test for equal variance of One-way ANOVA followed by Bonferroni post-hoc test for multiple comparison between the groups. Number of pairwise comparisons denoted are as follows; (a) NC vs $As_2O_3$(2mg) alone; (b) NC vs $As_2O_3$(4 mg) alone; (c) NC vs $As_2O_3$(8 mg) alone; (d) $As_2O_3$(2mg) alone vs $As_2O_3$(2mg)+CUR; (e) $As_2O_3$(4mg) alone vs $As_2O_3$(4mg)+CUR; (f) $As_2O_3$(8mg) alone vs $As_2O_3$(8mg)+CUR.

**S4:** Descriptive statistics of total ambulatory time (sec) of control and experimental groups during OFT.

| **Groups** | **Total Ambulatory time (sec)** | ***p* value** |
|---|---|---|
| Normal Control | 147.47 ± 44.03 | a) *p*< 1.000<br>b) *p*< .111<br>c) *p*< 1.000<br>d) *p*< 1.000<br>e) *p*< 1.000<br>f) *p*< 1.000 |
| Vehicle Control | 143 ± 53.19 | |
| Curcumin(100mg) | 142.08 ± 41.61 | |
| $As_2O_3$(2mg) | 146.27 ± 50.35 | |
| $As_2O_3$(4mg) | 88.92 ± 36.06 | |
| $As_2O_3$(8mg) | 138.76 ± 56.1 | |
| $As_2O_3$(2mg)+Cur(100mg) | 130.1 ± 56.33 | |
| $As_2O_3$(4mg)+Cur(100mg) | 116.37 ± 25.3 | |
| $As_2O_3$(8mg)+Cur(100mg) | 123.40 ± 53.01 | |
| **Overall ANOVA *p*< .0638** | | |

Data expressed as mean ± SD (n=12/group). Result analysed using Bartlett's test for equal variance of One-way ANOVA followed by Bonferroni post-hoc test for multiple comparison between the groups. Number of pairwise comparisons denoted are as follows; (a) NC vs $As_2O_3$(2mg) alone; (b) NC vs $As_2O_3$(4 mg) alone; (c) NC vs $As_2O_3$(8 mg) alone; (d) $As_2O_3$(2mg) alone vs $As_2O_3$(2mg)+CUR; (e) $As_2O_3$(4mg) alone vs $As_2O_3$(4mg)+CUR; (f) $As_2O_3$(8mg) alone vs $As_2O_3$(8mg)+CUR.

**S5:** Descriptive statistics of total distance travelled (metre) by control and experimental animals in OFT.

| Groups | Total Distance Travelled | *p* value |
|---|---|---|
| Normal Control | 8.3975 ± 3.19717 | |
| Vehicle Control | 7.528333 ± 3.621965 | |
| Curcumin(100mg) | 7.0975 ± 2.560554 | |
| $As_2O_3$(2mg) | 7.455833 ± 3.288599 | *a)* *p*< 1.000 |
| $As_2O_3$(4mg) | 5.311667 ± 2.301359 | *b)* *p*< .528 |
| $As_2O_3$(8mg) | 5.965 ± 2.430707 | *c)* *p*< 1.000 *d)* *p*< 1.000 |
| $As_2O_3$(2mg)+Cur(100mg) | 6.519167 ± 3.107377 | *e)* *p*< .747 *f)* *p*< 1.000 |
| $As_2O_3$(4mg)+Cur(100mg) | 8.885 ± 2.582372 | |
| $As_2O_3$(8mg)+Cur(100mg) | 7.436667 ± 3.886394 | |
| **Overall ANOVA *p*< .1233** | | |

Data expressed as mean ± SD (n=12/group). Result analysed using Bartlett's test for equal variance of One-way ANOVA followed by Bonferroni post-hoc test for multiple comparison between the groups. Number of pairwise comparisons denoted are as follows; (a) NC vs $As_2O_3$(2mg) alone; (b) NC vs $As_2O_3$(4 mg) alone; (c) NC vs $As_2O_3$(8 mg) alone; (d) $As_2O_3$(2mg) alone vs $As_2O_3$(2mg)+CUR; (e) $As_2O_3$(4mg) alone vs $As_2O_3$(4mg)+CUR; (f) $As_2O_3$(8mg) alone vs $As_2O_3$(8mg)+CUR.

**S6:** Descriptive statistics of time spent in centre, periphery and corner (sec) of the open field by control and experimental animals.

| Groups | Centre Time spent (A) | Periphery Time spent (B) | Corner Time spent (C) | *p* value (A,B,C) |
|---|---|---|---|---|
| Normal Control | 19.05 (9.55 - 53.1) | 114.51 ± 42.17 | 166.82 ± 41.66 | a) $p < .6033, 1.000, 1.000$<br>b) $p < .0998, .003, 1.000$<br>c) $p < .0016, .010, 1.000$<br>d) $p < .4884, 1.000, 1.000$<br>e) $p < .1123, 1.000, 1.000$<br>f) $p < .0832, .770, 1.000$ |
| Vehicle Control | 17.65(3.3 - 60.75) | 118.82 ± 47.4 | 155.27 ± 53.55 | |
| Curcumin(100mg) | 30.95(13.6 - 63.5) | 96.79 ± 32.55 | 166.85 ± 43.43 | |
| $As_2O_3$(2mg) | 18.9(7.05 - 53.9) | 112.29 ± 21.42 | 151.4 ± 45.63 | |
| $As_2O_3$(4mg) | 14.77(3.3 - 73.25) | 59.01 ± 26.43 | 187 ± 39.67 | |
| $As_2O_3$(8mg) | 9 (2.8 - 23) | 63.18 ± 35.54 | 189.77 ± 45.19 | |
| $As_2O_3$(2mg)+Cur(100mg) | 19.87(3.95 - 69.5) | 100.47 ± 34.11 | 171.16 ± 42.93 | |
| $As_2O_3$(4mg)+Cur(100mg) | 21.8(10.3 - 57.8) | 87.49 ± 28.35 | 187.42 ± 39.69 | |
| $As_2O_3$(8mg)+Cur(100mg) | 17.42 (4.05 - 88.05) | 95.04 ± 23.48 | 171.41 ± 40.93 | |
| **Overall ANOVA P value** | **$p < .0059$** | **$p < .0001$** | **$p < .3066$** | |

Data expressed as median (min-max)/mean ±SD (n=12/group). Results analysed using Bartlett's test for equal variance of One-way ANOVA followed by Bonferroni post-hoc test for multiple comparison between the groups. Number of pairwise comparisons denoted are as follows; (a) NC vs $As_2O_3$(2mg) alone; (b) NC vs $As_2O_3$(4 mg) alone; (c) NC vs $As_2O_3$(8 mg) alone; (d) $As_2O_3$(2mg) alone vs $As_2O_3$(2mg)+CUR; (e) $As_2O_3$(4mg) alone vs $As_2O_3$(4mg)+CUR; (f) $As_2O_3$(8mg) alone vs $As_2O_3$(8mg)+CUR.

**S7:** Descriptive statistics of centre entries, peripheral entries, corner entries (n) of control and experimental groups in OFT.

| **Groups** | **Centre Entries (A)** | **Periphery Entries (B)** | **Corner Entries (C)** | ***p* value (A,B,C)** |
|---|---|---|---|---|
| Normal Control | 6.75(1.5 - 17.5) | 34.75 (13 - 55.5) | 26 ± 9.58 | a) $p < .68533$, 1.000, 1.000<br>b) $p < .0668$, .811, 1.000<br>c) $p < .0034$, 1.000, 1.000<br>d) $p < .4180$, 1.000, 1.000<br>e) $p < .6226$, 1.000, .169<br>f) $p < .1245$, 1.000, .001 |
| Vehicle Control | 7.75(1.5-16.5) | 29.75(11 - 58) | 21.66 ± 8.61 | |
| Curcumin(100mg) | 8.25(3 - 15) | 30.75(15.5 - 50.5) | 21 ± 6.59 | |
| $As_2O_3$(2mg) | 7.5(1 - 13) | 30.75(11 - 47.5) | 21.62 ± 9.37 | |
| $As_2O_3$(4mg) | 4.25(1 - 7.5) | 21.5(1 - 34) | 24.62 ± 8.02 | |
| $As_2O_3$(8mg) | 3 (1 - 4.5) | 22 (13 - 49.5) | 27.7 ± 11.75 | |
| $As_2O_3$(2mg)+Cur(100mg) | 4.75(2 - 13.5) | 23.75(10 - 50) | 19.5 ± 8.59 | |
| $As_2O_3$(4mg)+Cur(100mg) | 4.5(1 - 12) | 24(15 - 43) | 14.79 ± 5.51 | |
| $As_2O_3$(8mg)+Cur(100mg) | 4.5(1 - 9) | 27.25(10 - 54) | 12.16 ± 4.46 | |
| **Overall ANOVA p-value** | ***p*< .0013** | ***p*< .490** | ***p*< .0001** | |

Data expressed as median (min-max)/mean ±SD (n=12/group). Results analysed using Bartlett's test for equal variance of One-way ANOVA followed by Bonferroni post-hoc test for multiple comparison between the groups. Number of pairwise comparisons denoted are as follows; (a) NC vs $As_2O_3$(2mg) alone; (b) NC vs $As_2O_3$(4 mg) alone; (c) NC vs $As_2O_3$(8 mg) alone; (d) $As_2O_3$(2mg) alone vs $As_2O_3$(2mg)+CUR; (e) $As_2O_3$(4mg) alone vs $As_2O_3$(4mg)+CUR; (f) $As_2O_3$(8mg) alone vs $As_2O_3$(8mg)+CUR.

.

**S8:** Descriptive statistics of recognition index & discrimination index of control and experimental groups in NORT.

| **Groups** | **Recognition Index (RI)** | **Discrimination Index (DI)** |
|---|---|---|
| Normal Control | 0.66(0 - 1) | 0.33(-0.86 - 1) |
| Vehicle Control | 0.63(0 - 1) | 0.26(-0.41 - 1) |
| Curcumin(100mg) | 0.55(0.1 - 1) | 0.37(-0.33 - 1) |
| $As_2O_3$(2mg) | 0.59(0 - 0.95) | 0.18(-0.92 - 0.91) |
| $As_2O_3$(4mg) | 0.38(0 -0.75) | 0(-1 - 0.5) |
| $As_2O_3$(8mg) | 0.33(0 - 0.92) | -0.08(-1 - 1) |
| $As_2O_3$(2mg)+Cur(100mg) | 0.5(0.32 - 0.92) | 0.1(-0.79 - 1) |
| $As_2O_3$(4mg)+Cur(100mg) | 0.56(0 - 1) | 0(-0.6 - 1) |
| $As_2O_3$(8mg)+Cur(100mg) | 0.36(0 - 1) | 0.01(-1 - 1) |
| **Overall ANOVA p value** | ***p*<.2595** | ***p*< .2696** |

Data expressed as median (min-max) (n=12/group). Results analysed using Bartlett's test for equal variance of One-way ANOVA followed by Bonferroni post-hoc test for multiple comparison between the groups. Number of pairwise comparisons denoted are as follows; (a) NC vs $As_2O_3$(2mg) alone; (b) NC vs $As_2O_3$(4 mg) alone; (c) NC vs $As_2O_3$(8 mg) alone; (d) $As_2O_3$(2mg) alone vs $As_2O_3$(2mg)+CUR; (e) $As_2O_3$(4mg) alone vs $As_2O_3$(4mg)+CUR; (f) $As_2O_3$(8mg) alone vs $As_2O_3$(8mg)+CUR.

**S9:** Descriptive statistics of latency to fall (sec) in control and experimental groups during rota-rod test.

| **Groups** | **Latency to fall (sec)** | ***p* Value** |
|---|---|---|
| Normal Control | 36.23(20.2 - 64.33) | a) $p < .0079$<br>b) $p < .0015$<br>c) $p < .0005$<br>d) $p < .1333$<br>e) $p < .0327$<br>f) $p < .0496$ |
| Vehicle Control | 36.88(14.9 - 69.37) | |
| Curcumin(100mg) | 28.88(10.9 - 60) | |
| $As_2O_3$(2mg) | 17.01 (2.5 - 39.6) | |
| $As_2O_3$(4mg) | 10.25(2.37 - 37.2 ) | |
| $As_2O_3$(8mg) | 11.76 (2.27 - 37.97 ) | |
| $As_2O_3$(2mg)+Cur(100mg) | 21.65(10.9 - 66.3) | |
| $As_2O_3$(4mg)+Cur(100mg) | 24.81 (12.5 - 38.47) | |
| $As_2O_3$(8mg)+Cur(100mg) | 19(11.27 - 37.2) | |
| **Overall ANOVA $p < .0001$** | | |

Data expressed as median (min-max) (n=12/group). Results analysed using Bartlett's test for equal variance of One-way ANOVA followed by Bonferroni post-hoc test for multiple comparison between the groups. Number of pairwise comparisons denoted are as follows; (a) NC vs $As_2O_3$(2mg) alone; (b) NC vs $As_2O_3$(4 mg) alone; (c) NC vs $As_2O_3$(8 mg) alone; (d) $As_2O_3$(2mg) alone vs $As_2O_3$(2mg)+CUR; (e) $As_2O_3$(4mg) alone vs $As_2O_3$(4mg)+CUR; (f) $As_2O_3$(8mg) alone vs $As_2O_3$(8mg)+CUR.

**S10:** Descriptive statistics of average escape latency (sec) during acquisition period of control and experimental groups in MWM.

| Average Escape latency (sec) during acquisition days | | | | |
|---|---|---|---|---|
| **Groups** | **Day 1** | **Day 2** | **Day 3** | **Day 4** |
| Normal Control | 41.45 ( 15.9 - 60 ) | 25.35( 10.02 - 60 ) | 27.68( 7.15 - 60 ) | 24.31( 4.3 - 42.05 ) |
| Vehicle Control | 23.37( 3.72 - 58.3 ) | 19.92( 3.82 - 35.12 ) | 12.7( 5.05 - 51.52 ) | 6.91( 2.3 - 41.02 ) |
| Curcumin(100mg) | 34.06( 4.1 - 45.27 ) | 11.16( 4.42 - 60 ) | 22.36( 3.37 - 47.37 ) | 11.51( 3.7 - 31.62 ) |
| $As_2O_3$(2mg) | 33.51( 4.47 - 60 ) | 21.5( 6.45 - 60 ) | 18( 4.62 - 60 ) | 15.53( 3.45 - 60 ) |
| $As_2O_3$(4mg) | 39.81( 11.3 - 60 ) | 26.18( 12.3 - 60 ) | 25.64( 13.47 - 60 ) | 37.19( 13.17 - 60 ) |
| $As_2O_3$(8mg) | 28.78( 11.92 - 60 ) | 28.02( 17.07 - 60 ) | 23.74( 4.22 - 60 ) | 23.58( 11.4 - 60 ) |
| $As_2O_3$(2mg)+Cur(100mg) | 34.01( 10.67 - 60 ) | 19.68( 5.47 - 60 ) | 17.23( 3.4 - 46.92 ) | 15.73( 4.42 - 39.57 ) |
| $As_2O_3$(4mg)+Cur(100mg) | 35.81( 18.22 - 60 ) | 36.31( 10.6 - 60 ) | 27.2( 9.72 - 60 ) | 22.02( 3.82 - 43.45 ) |
| $As_2O_3$(8mg)+Cur(100mg) | 35.37( 15.97 - 60 ) | 33.97( 12.6 - 60 ) | 22.86( 9.5 - 50.15 ) | 21.15( 5.62 - 60 ) |
| **Overall ANOVA *p* value** | ***p*< .4441** | ***p*< .0784** | ***p*< .2208** | ***p*< .0006** |

**S11:** Descriptive statistics of average distance travelled (m) during acquisition phase of control and experimental groups in MWM test.

| **Average Distance travelled during acquisition days** | | | | |
|---|---|---|---|---|
| **Groups** | **Day 1** | **Day 2** | **Day 3** | **Day 4** |
| Normal Control | 4.44 (1.59 - 6.72) | 4.18(2.64 - 9.09) | 3.76(1.34 - 6.12) | 3.48(0.9 - 7.19) |
| Vehicle Control | 3.25(0.66 - 6.71) | 2.26(0.86 - 4.92) | 2.31(0.76 - 5.03) | 1.22(0.44 - 6.55) |
| Curcumin(100mg) | 4.77(0.64 - 7.78) | 2.16(0.97 - 7.23) | 3.39(1.29 - 6.6) | 2.58(0.66 - 6.26) |
| $As_2O_3$(2mg) | 4.62(1.28 - 8.08) | 3.59(1.17 - 7.27) | 3.27(0.99 - 6.28) | 2.37(0.78 - 6.57) |
| $As_2O_3$(4mg) | 4.32 (2.23 - 8.08) | 3.71(2.32 - 7.27) | 3.45(1.91 - 5.47) | 3.56(2.57 - 6.57) |
| $As_2O_3$(8mg) | 4.58 (2.04 - 5.58) | 3.9 (2.99 - 5.72) | 3.38 (0.77 - 7.51) | 3.84(1.97 - 6.64) |
| $As_2O_3$(2mg)+Cur(100mg) | 4.45(2.23 - 6.62) | 2.81(1.11 - 4.58) | 2.34(0.68 - 5.57) | 2.57(0.8 - 3.83) |
| $As_2O_3$(4mg)+Cur(100mg) | 5.27(2.61 - 8.51) | 4.29(2.03 - 7.27) | 4.79(2.11 - 7.36) | 4.3(0.8 - 8.23) |
| $As_2O_3$(8mg)+Cur(100mg) | 4.97(1.46 - 8.08) | 4.25(1.04 - 7.27) | 3.39(2 - 6.28) | 3.19(1.24 - 6.57) |
| **Overall ANOVA** <br> ***p* value** | ***p*<.5511** | ***p*< .0174** | ***p*< .0856** | ***p*< .0017** |

**S12:** Descriptive statistics of average swim speed (m/sec) during acquisition period of control and experimental groups in MWM.

| **Average swim speed (m/sec) during acquisition days** | | | | |
|---|---|---|---|---|
| **Groups** | **Day 1** | **Day 2** | **Day 3** | **Day 4** |
| Normal Control | 0.09 ± 0.02 | 0.09 ± 0.02 | 0.08 ± 0.01 | 0.09 ± 0.02 |
| Vehicle Control | 0.08 ± 0.02 | 0.07 ± 0.02 | 0.07 ± 0.02 | 0.06 ± 0.02 |
| Curcumin(100mg) | 0.09 ± 0.03 | 0.08 ± 0.02 | 0.09 ± 0.02 | 0.08 ± 0.02 |
| $As_2O_3$(2mg) | 0.09 ± 0.02 | 0.08 ± 0.02 | 0.09 ± 0.01 | 0.07 ± 0.02 |
| $As_2O_3$(4mg) | 0.09 ± 0.01 | 0.09 ± 0.02 | 0.08 ± 0.01 | 0.08 ± 0.02 |
| $As_2O_3$(8mg) | 0.09 ± 0.01 | 0.09 ± 0.01 | 0.08 ± 0.01 | 0.08 ± 0.01 |
| $As_2O_3$(2mg)+Cur(100mg) | 0.08 ± 0.02 | 0.07 ± 0.01 | 0.07 ± 0.02 | 0.07 ± 0.01 |
| $As_2O_3$(4mg)+Cur(100mg) | 0.11 ± 0.03 | 0.09 ± 0.03 | 0.1 ± 0.02 | 0.11 ± 0.03 |
| $As_2O_3$(8mg)+Cur(100mg) | 0.09 ± 0.03 | 0.09 ± 0.03 | 0.09 ± 0.01 | 0.08 ± 0.02 |
| **Overall ANOVA *p* value** | ***p*<.026** | ***p*<.364** | ***p*<.327** | ***p*<.317** |

**S13:** Descriptive statistics showing number of platform crossing (n) and time (sec) spent in platform quadrant by control and experimental animals during probe trial of MWM test.

| **Groups** | **No. of Platform crossing (A)** | **Time Spent in Platform Quadrant (B)** | ***p* Value (A,B)** |
|---|---|---|---|
| Normal Control | 2(1 - 5) | 17.65(9.4 - 22) | a) *p*< .1002, .0208<br>b) *p*< .0665, .1057<br>c) *p*< .1628, .1655<br>d) *p*< .6990, .0991<br>e) *p*< .0422, .0410<br>f) *p*< .0363, .0433 |
| Vehicle Control | 3(1 - 7) | 20.65(10.1 - 32.3) | |
| Curcumin(100mg) | 2(0 - 4) | 14.6(10.5 - 23.8) | |
| $As_2O_3$(2mg) | 1.5(0 - 4) | 11.5(0 - 33.3) | |
| $As_2O_3$(4mg) | 1(0 - 5) | 13.1(0 - 27.7) | |
| $As_2O_3$(8mg) | 2(1 - 3) | 12.85(9.2 - 22.9) | |
| $As_2O_3$(2mg)+Cur(100mg) | 1.5(0 - 5) | 7.95(0 - 41.6) | |
| $As_2O_3$(4mg)+Cur(100mg) | 3(0 - 4) | 16 (12.6 - 33.3) | |
| $As_2O_3$(8mg)+Cur(100mg) | 2.5(1 - 3) | 21.3(0.8 - 36.4) | |
| **Overall ANOVA *p* value** | ***p*< .0486** | ***p*< .0002** | |

Data expressed as median (min-max) (n=12/group). Results analysed using Bartlett's test for equal variance of One-way ANOVA followed by Bonferroni post-hoc test for multiple comparison between the groups. Number of pairwise comparisons denoted are as follows; (a) NC vs $As_2O_3$(2mg) alone; (b) NC vs $As_2O_3$(4 mg) alone; (c) NC vs $As_2O_3$(8 mg) alone; (d) $As_2O_3$(2mg) alone vs $As_2O_3$(2mg)+CUR; (e) $As_2O_3$(4mg) alone vs $As_2O_3$(4mg)+CUR; (f) $As_2O_3$(8mg) alone vs $As_2O_3$(8mg)+CUR.

**S14:** Descriptive statistics of status of antioxidant enzymes (GSH, GPx) and oxidative stress marker (MDA) across control and experimental groups.

| **Groups** | **GSH (A)** | **GPx (B)** | **MDA (C)** | ***p* Value (A,B,C)** |
|---|---|---|---|---|
| Normal Control | 4.62 (2.15 - 7.2) | 32.50 ± 4.73 | 961.85 ± 118.68 | a) *p*< .0247, 1.000, .072<br>b) *p*< .0104, .270, 1.000<br>c) *p*< .0038, .488, .009<br>d) *p*< .8726, 1.000, .302<br>e) *p*< .0547, .889, .348<br>f) *p*< .0038, 1.000, .005 |
| Vehicle Control | 3.71(1.19 - 5.68) | 30.25 ± 6.12 | 847.69 ± 198.75 | |
| Curcumin(100mg) | 3.44(2.11 - 4.81) | 31.98 ± 10.67 | 900.16 ± 185.8 | |
| $As_2O_3$(2mg) | 1.67(0.91 - 4.63) | 25.04 ± 3.16 | 1478.84 ± 129.62 | |
| $As_2O_3$(4mg) | 1.81(0.18 - 3.59) | 23.08 ± 4.44 | 1285.41 ± 284.44 | |
| $As_2O_3$(8mg) | 0.55(0.32 - 0.96) | 23.84 ± 5.86 | 1587.34 ± 238.15 | |
| $As_2O_3$(2mg)+Cur(100mg) | 2.06 (1.1 - 4.58) | 29.20 ± 7.10 | 1044.55 ± 481.78 | |
| $As_2O_3$(4mg)+Cur(100mg) | 2.95(2.52 - 3.57) | 30.91 ± 3.54 | 859.61 ± 327.33 | |
| $As_2O_3$(8mg)+Cur(100mg) | 4.71(1 - 5.55) | 30.67 ± 7.86 | 926.76 ± 297.05 | |
| **Overall ANOVA *p* value** | ***p*< .0009** | ***p*< .0346** | ***p*< .001** | |

Data expressed as median (min-max)/mean±SD (n=7/group). Results analysed using Bartlett's test for equal variance of One-way ANOVA followed by Bonferroni post-hoc test for multiple comparison between the groups. Number of pairwise comparisons denoted are as follows; (a) NC vs $As_2O_3$(2mg) alone; (b) NC vs $As_2O_3$(4 mg) alone; (c) NC vs $As_2O_3$(8 mg) alone; (d) $As_2O_3$(2mg) alone vs $As_2O_3$(2mg)+CUR; (e) $As_2O_3$(4mg) alone vs $As_2O_3$(4mg)+CUR; (f) $As_2O_3$(8mg) alone vs $As_2O_3$(8mg)+CUR.

**S15:** Observations of morphometric parameter- neuronal area (µm$^2$) in dorsal & ventral striatum of control and experimental groups.

| Groups | Area of neuron in dorsal striatum (A) | Area of neuron in ventral striatum (B) | *p* value (A,B) |
|---|---|---|---|
| Normal Control | 52.96 ± 6.87 | 55.79 ± 13.09 | a) *p*< .676, 1.000<br>b) *p*< 1.000, 1.000<br>c) *p*< 1.000, 1.000<br>d) *p*< 1.000, 1.000<br>e) *p*< 1.000, 1.000<br>f) *p*< 1.000, 1.000 |
| Vehicle Control | 53.6 ± 3.26 | 54.99 ± 0.88 | |
| Curcumin(100mg) | 60.28 ± 8.51 | 68.12 ± 7.88 | |
| $As_2O_3$(2mg) | 63.95 ± 10.86 | 65.31 ± 9.8 | |
| $As_2O_3$(4mg) | 51.57 ± 6.34 | 49.95 ± 3.07 | |
| $As_2O_3$(8mg) | 51.71 ± 6.43 | 52.52 ± 5.32 | |
| $As_2O_3$(2mg)+Cur(100mg) | 59.45 ± 5.08 | 64.93 ± 4.82 | |
| $As_2O_3$(4mg)+Cur(100mg) | 59.78 ± 8.64 | 59.25 ± 12.1 | |
| $As_2O_3$(8mg)+Cur(100mg) | 61 ± 10.9 | 59.29 ± 3.36 | |
| **Overall ANOVA p value** | ***p*< .0513** | ***p*< .0018** | |

Data expressed as mean±SD (n=6/group). Results analysed using Bartlett's test for equal variance of One-way ANOVA followed by Bonferroni post-hoc test for multiple comparison between the groups. Number of pairwise comparisons denoted are as follows; (a) NC vs $As_2O_3$(2mg) alone; (b) NC vs $As_2O_3$(4 mg) alone; (c) NC vs $As_2O_3$(8 mg) alone; (d) $As_2O_3$(2mg) alone vs $As_2O_3$(2mg)+CUR; (e) $As_2O_3$(4mg) alone vs $As_2O_3$(4mg)+CUR; (f) $As_2O_3$(8mg) alone vs $As_2O_3$(8mg)+CUR.

**S16:** Observations of morphometric parameters- neuron density (cells/µm$^2$) in dorsal and ventral striatum of control and experimental animals.

| Groups | Neuron density in dorsal striatum (A) | Neuron density in ventral striatum (B) | p value (A,B) |
|---|---|---|---|
| Normal Control | 63.12(13.75 - 124) | 50.87 (13.25 - 104.75) | a) *p*< .7488, .8728<br>b) *p*< 1.000, 1.000<br>c) *p*< 1.000, 1.000<br>d) *p*< .1488, .2002<br>e) *p*< .4225, .3776<br>f) *p*< .4225, .3776 |
| $As_2O_3$(2mg) | 54.62(21.5 - 59.5) | 42.25(22.25 - 58.25) | |
| $As_2O_3$(4mg) | 54.87(43.75 - 57.5) | 54.87(37 - 59.75) | |
| $As_2O_3$(8mg) | 54.87(43.75 - 57.5) | 54.87(37 - 59.75) | |
| $As_2O_3$(2mg)+Cur(100mg) | 58.75(52 - 61) | 48.5(45.75 - 57.25) | |
| $As_2O_3$(4mg)+Cur(100mg) | 50.62(28.5 - 77.75) | 46.62(37 - 97.25) | |
| $As_2O_3$(8mg)+Cur(100mg) | 50.62(28.5 - 77.75) | 46.62(37 - 97.25) | |
| **Overall ANOVA p value** | ***p*< .0153** | ***p*< .0051** | |

Data expressed as median (min-max) (n=6/group). Results analysed using Bartlett's test for equal variance of One-way ANOVA followed by Bonferroni post-hoc test for multiple comparison between the groups. Number of pairwise comparisons denoted are as follows; (a) NC vs $As_2O_3$(2mg) alone; (b) NC vs $As_2O_3$(4 mg) alone; (c) NC vs $As_2O_3$(8 mg) alone; (d) $As_2O_3$(2mg) alone vs $As_2O_3$(2mg)+CUR; (e) $As_2O_3$(4mg) alone vs $As_2O_3$(4mg)+CUR; (f) $As_2O_3$(8mg) alone vs $As_2O_3$(8mg)+CUR.
